# Integrative 4D-conformational mechanisms of single AdiC transporter molecules

**DOI:** 10.1101/2022.07.29.501984

**Authors:** John H. Lewis, Yufeng Zhou, Zhe Lu

## Abstract

To understand the mechanism of counter-transport of substrates by the amino-acid transporter AdiC, we used a state-of-the-art polarization-microscope to investigate conformation-specific changes of the emission polarization of a fluorophore attached to individual AdiC molecules. This capability enabled us to determine the lifetimes of two energetic states of each of AdiC’s four conformations in the absence and presence of its two natural substrates, totaling 24 states. From these lifetimes and relative state-to-state transition frequencies, we further determined 60 rate constants of all state transitions and the 4 K_D_ values for the two substrates to interact with both sides of AdiC, quantitatively defining a 24-state model that satisfactorily predicts previously observed transporting behaviors of AdiC. Combining this temporal information and the existing structural information, we have successfully built a fully experiment-based integrative 4D-model to capture and exhibit the complex spatiotemporal mechanisms of a facilitated counter-transport of an amino acid and its metabolite. Thus, a combination of the present method and existing structural techniques serves as an effective means to help transition structural biology, which has thus far been highly successful in the investigation of individual static structures, to an integrative form of dynamic structural biology.

## Introduction

Biological transporters comprise a very large collection of proteins that enable specific impermeable substances to get across cell membranes along or against a concentration gradient though a series of complex conformational changes. Based on kinetic properties of transporter function, four basic types of conformation were conceived in terms of the accessibility of its extracellular and intracellular sides to substances, namely, being externally open (E_o_), externally occluded (E_x_), internally open (I_o_), and internally occluded (I_x_) states, where the unspecified side of the transporter is always inaccessible (Post et al., 1972, Shi, 2013, Krammer and Prevost, 2019). Structural biology studies have revealed the detailed structural features underlying these functionally defined conformations for numerous transporters (Shi, 2013, Krammer and Prevost, 2019).

Despite the tremendous progress made in functional and structural studies of transporters, some important and technically challenging questions remain. Among them, because protein-conformational changes are defined in terms of both spatial and temporal characteristics, it is important to know how the individual structural states are temporally related. To accomplish this task, the required temporal measurements at each time point need to contain the spatial information that can unambiguously identify the specific conformational state adopted by the protein. With relatable temporal information and structural information, one could generate an integrative mechanistic model in all four dimensions (4D) to account for the spatiotemporal behaviors of a protein molecule.

Furthermore, like the kinetics of an enzyme (Plowman, 1971), those of an individual transporter is typically characterized functionally by its substrate turnover rate (*k*_cat_), and the substrate concentration (*K*_m_) required for achieving half of *k*_cat_. It is challenging to understand these two empirical function-defining parameters in terms of a kinetic mechanism involving the four types of basic conformation. Furthermore, each type of conformation may have more than one energetic state, and each state may be in an apo or substrate bound form. Thus, what underlies both *k*_cat_ and *K*_m_ are the rate constants that govern the transitions among an often very large number of states. To address this and the above mechanistic questions require the capability to quantitatively track the rapid conformational changes at the single-molecule level, which often occur on an angstrom scale.

In the companion manuscript, we demonstrate our successful resolution of changes among four conformations of single AdiC transporter molecules through monitoring the polarization changes of an attached fluorophore with a polarization microscope system assembled in house (Lewis and Lu, 2019c, Lewis and Lu, 2019a, Lewis and Lu, 2019b). Naturally, this transport facilitates both the uptake of the amino acid arginine (Arg^+^) into enterobacteria from the aqueous environment and the discharge of agmatine (Agm^2+^) enzymatically generated from Arg^+^ inside bacteria, thereby acting as a key component of a proton-extrusion system (Gong et al., 2003, Foster, 2004, Fang et al., 2007, Iyer et al., 2003, Krammer and Prevost, 2019). This system is critical for common pathogenic enterobacteria to pass the highly acidic gastric environment to reach the intestines.

Here, we set out to investigate the lifetimes of the conformational states, thereby determining the energetic states for each type of structural conformation and, furthermore, the relative state-to-state transition frequencies, thereby determining rate constants for the transitions among all identified states. With this collection of acquired information, we will establish a quantitative, mechanistic model to account for the functional characteristics of the transporter and, furthermore, will combine the acquired temporal information and available structural information to build an integrative 4D model to capture and illustrate our understanding of the transporter’s spatiotemporal mechanisms.

## Results

### Recordings of fluorescence intensity and determination of fluorophore orientation

Fig. 1A shows a set of intensity traces recorded at a rate of 10 ms per frame from a bifunctional rhodamine attached to helix-6A of AdiC, which was inserted into nanodiscs, attached to a coverslip, and exposed to 50 μM of the ligand Agm^2+^. Judging from crystal structures, helix-6A adopts different orientations in different states (Gao et al., 2009, Fang et al., 2009, Gao et al., 2010, Kowalczyk et al., 2011, Ilgu et al., 2016, Ilgu et al., 2021). Unless specified otherwise, all references to AdiC, a structural dimer, are in the context of an independent functional subunit; Agm^2+^ or Arg^+^ are referred to as substrate and ligand in the context of their transport and binding, respectively.

**Figure 1.**
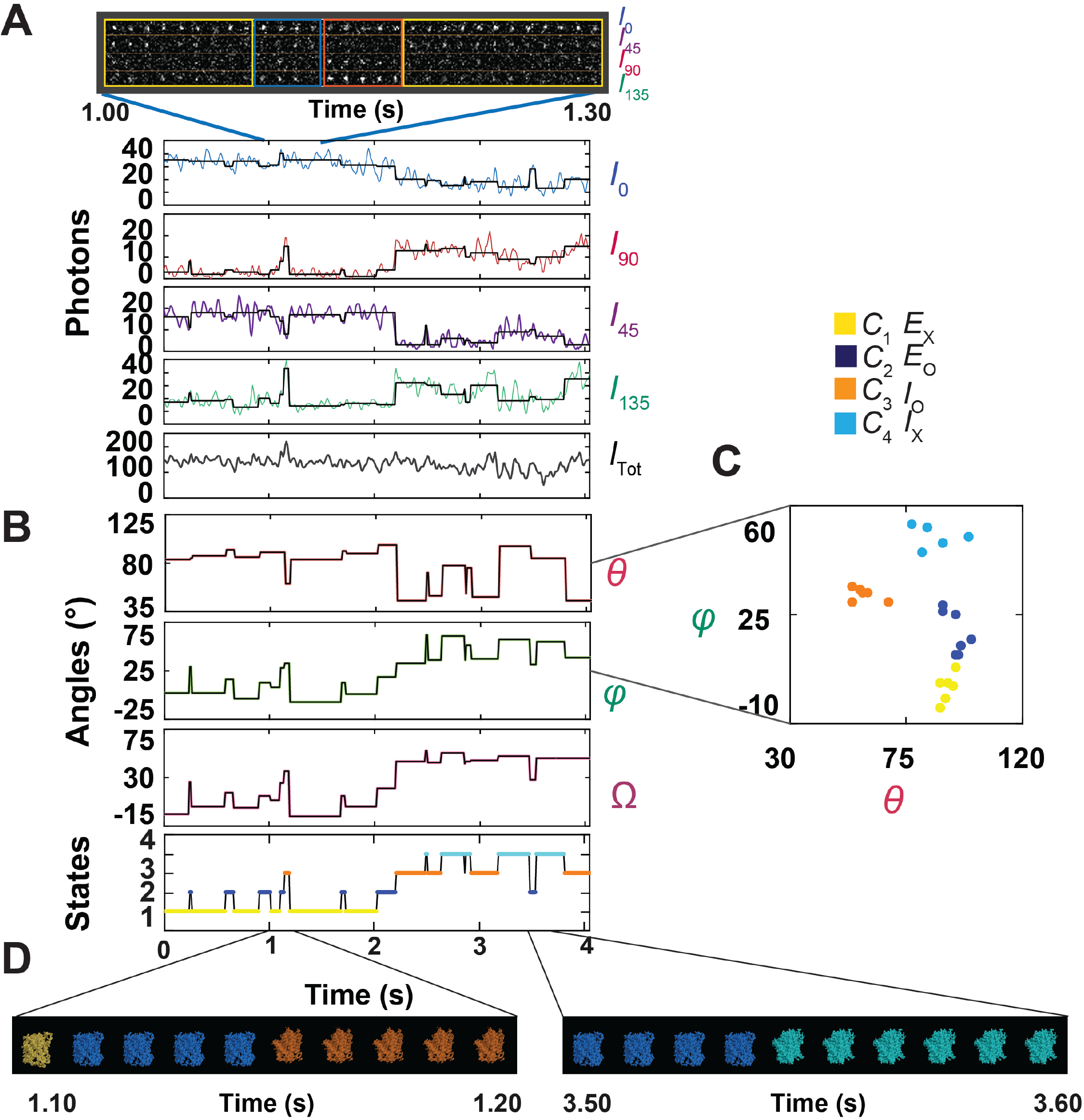
Polarized intensity components of a single fluorescent particle, *θ* and *φ* angles calculated from these components, and segments of a movie of conformational changes. (A) A segment of consecutive frames of four intensity components (*I*_0_, *I*_45_, *I*_90_ and *I*_135_) of a bifunctional-rhodamine-labeled apo AdiC molecule attached to an AdiC molecule obtained in the presence of 50 μM Agm^2+^. The integrated intensity values, color-coded for these intensities, and calculated *I*_tot_ values, are plotted against the observation time. Each vertical line in the black traces, superimposed on the colored intensity traces, indicates the time point at which a change in the fluorophore’s orientation is identified, whereas each horizontal line represents the mean intensity between two identified consecutive time points. (B) Traces for *θ* and *φ* in a local frame of reference are calculated from black intensity traces in A. Values of *Ω* are calculated relative to *C*_1_ from the changes in *θ* and *φ*. The bottom trace illustrates the state profile. (C) A scatter plot of *θ* versus *φ* determined from the traces shown in B. D. Consecutive frames in two segments of a movie of AdiC conformational changes where the four conformations are represented by the corresponding electron density maps (PDB: 7O82, 3L1L, 3GIA, 6F2G), in accordance with the temporal information encoded in the state-profile trace at bottom of B.

As shown in the companion manuscript, the recorded intensities were sorted into four polarized components (*I*_0_, *I*_45_, *I*_90_ and *I*_135_), from which we calculated the total intensity *I*_tot_ (Fig. 1A). The vertical black lines superimposed on the recorded intensities indicate the individual time points at which changes in the detected number of photons due to alterations in the fluorophore’s orientation occurred. Each horizontal line between a pair of consecutive vertical lines superimposed over the intensity traces corresponds to the average intensity value over the defined period. After acquiring this temporal information, averaging the intensities over the time that the AdiC molecule adopted a particular state, dubbed dwell time, markedly increased the signal-to-noise ratio (SNR) and thus angle resolution. From the resulting black traces, we calculated the angles *θ* and *φ* (expressed in a local framework of AdiC) for a given state, and then the direct angle change *Ω* between two states from their *θ* and *φ* values (Fig. 1B). For each ligand condition, we have identified four conformational state populations (*C*_1_-*C*_4_) from the values of *θ* and *φ* together, relating *C*_1_ to the externally occluded (E_x_), *C*_2_ to the externally open (E_o_), *C*_3_ to the internally open (I_o_), and *C*_4_ to the internally occluded (I_x_) states (Fig. 1C).

With the above information, we can generate the temporal template for Supplementary Video 1 to illustrate the conformational changes by matching the states identified at each time point to their corresponding atomic structures in the form of electron density maps, without any kinetic or structural modeling. This template was obtained from the state transition and identity information encoded in the *Ω* trace (Fig. 1B), under which is an illustrative color-coded state profile. The conformational states are shown in the form of electron density maps (PDB: 7O82, 3L1L, 3GIA, 6F2G) (Shaffer et al., 2009, Gao et al., 2010, Errasti-Murugarren et al., 2019, Ilgu et al., 2021), colored for states as for the state profile trace. Two segments of video frames from the movie are shown in Fig. 1D. Note that due to the limit of allowable file size, the video was made to be viewed in a small window of such software as Windows Media Player.

### Construction and analysis of dwell-time distributions of conformational states

Detection of transition points and identification of conformational states enabled us to measure the dwell times of an AdiC molecule in individual conformational states. Using *C*_2_ (E_o_) and *C*_3_ (I_o_) as examples, we show in Fig. 2 the distributions of dwell times in the absence or the presence of a saturating concentration of Arg^+^ or Agm^2+^. To analyze the dwell-time distributions, we used an exponential function derived for data recorded with a camera at a constant frame rate (Lewis et al., 2017) (Eq. 15; equations with a number greater than 14 are in Methods), which also addresses the problem arising from missing short events. As shown in Fig. 2, for all conformational states under all examined conditions, dwell-time distributions were statistically much better fitted with a double exponential function than a single exponential function (Eq. 16), based on all p-values being less than 10^−16^ in F-tests (Woody et al., 2016). Thus, each of the four conformational states has two identifiable, energetically distinct, and kinetically connected states. Consequently, 8 states are required to account for the behaviors of AdiC in the absence or the presence of either ligand type, totaling 24 states (Fig. 3). To reflect this expansion, the *C*_1_ conformation is split to two energetic states *S*_1_ and *S*_5_, *C*_2_ to *S*_2_ and *S*_6_, *C*_3_ to *S*_3_ and *S*_7_, and *C*_4_ to *S*_4_ and *S*_8_. Regarding ligand accessibility, among all states only *S*_6_ and *S*_7_ are accessible to the internal and the external ligand, respectively. *S*_5_ and *S*_8_ are treated as occluded *“cul-de-sac”* states that cannot directly transition to any open states.

**Figure 2.**
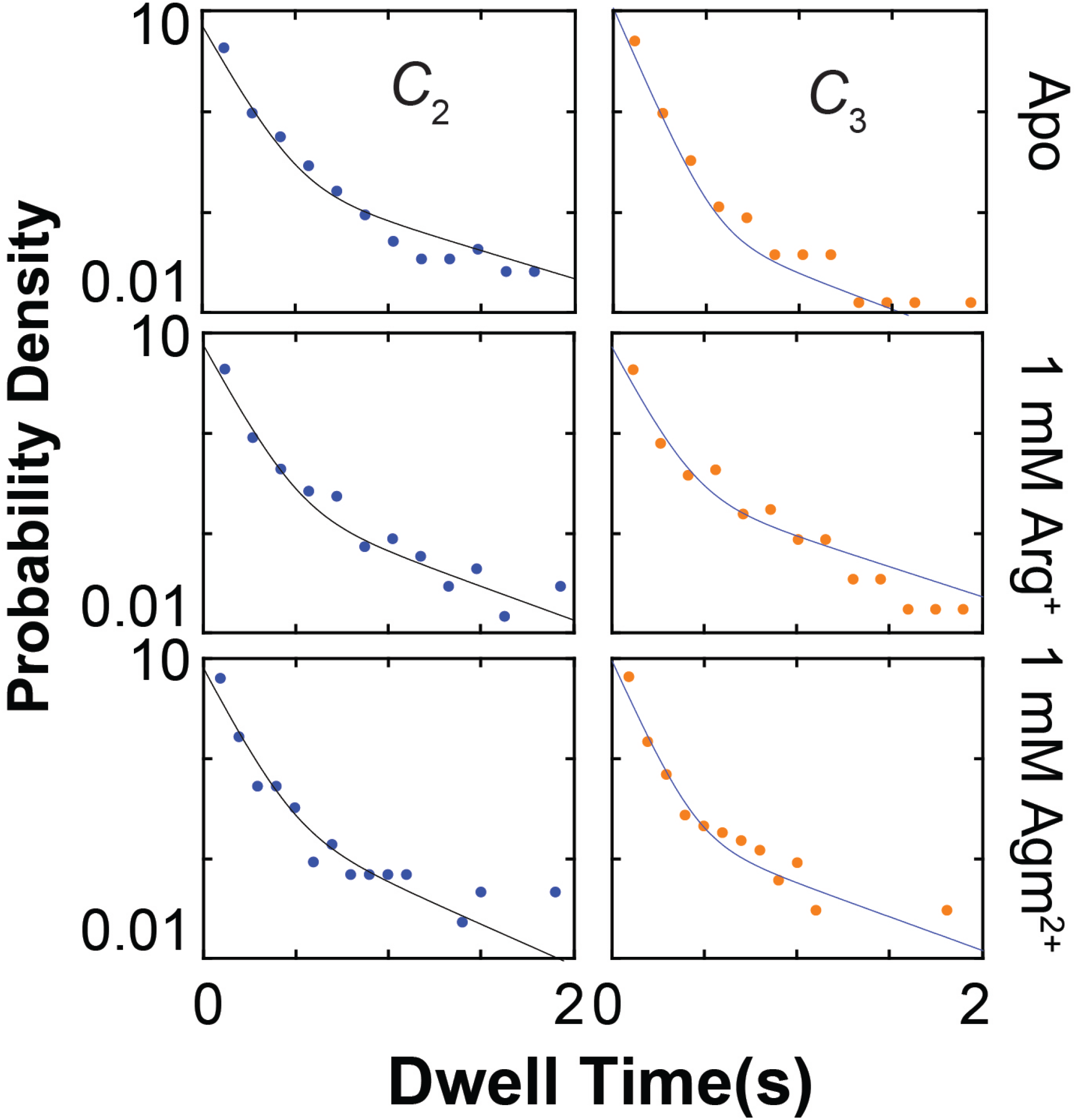
Examples of dwell time distributions of conformational states. The distributions for *C*_2_ (left) and *C*_3_ (right) in the absence (top) or presence of 1 mM Arg^+^ (middle) or 1 mM Agm^2+^ (bottom), plotted as the probability density on a log scale against dwell time of a given state, all exhibit two distinct phases. Each distribution was built on the basis of ~2000-5700 dwelling events in individual states of ~100-180 AdiC molecules. The curves superimposed on the data are fits of a two-exponential equation (Eq. 15).

**Figure 3.**
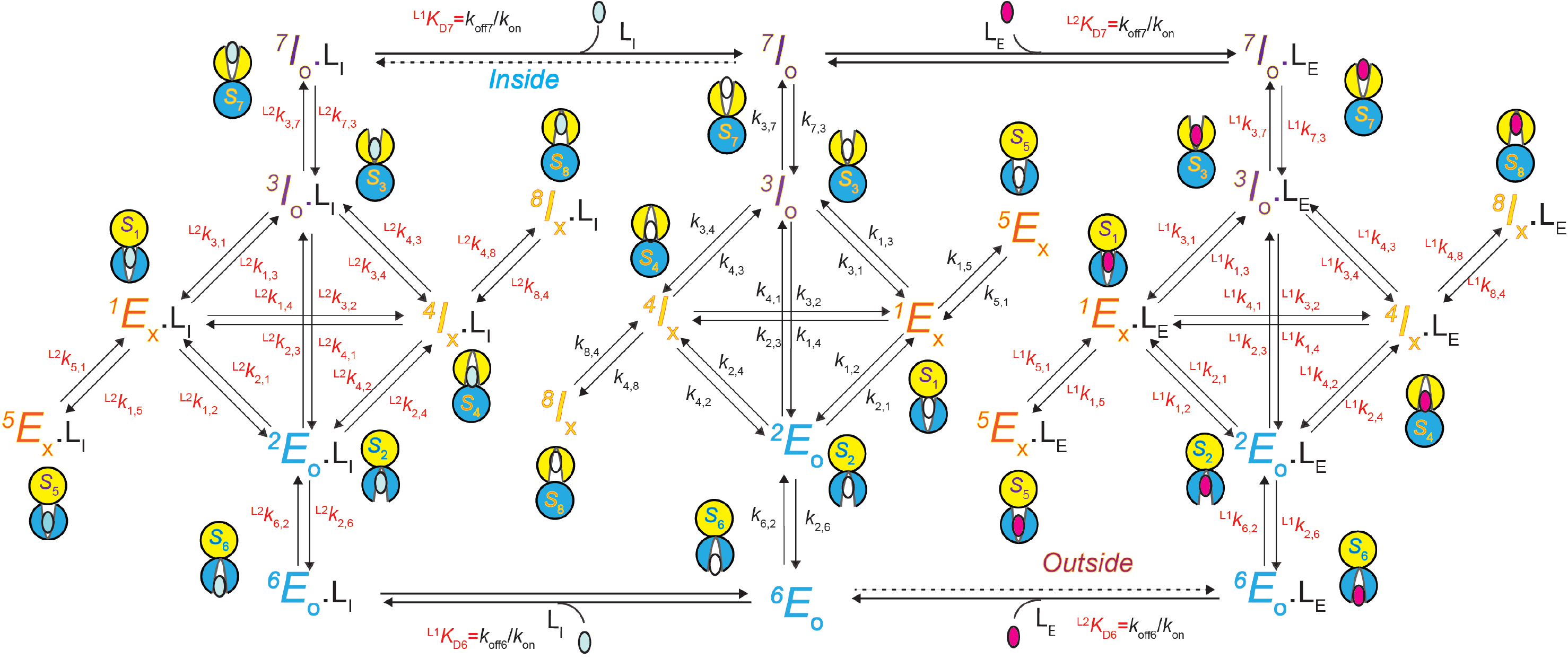
A 24-state kinetic model of AdiC for transporting two types of ligand on opposite sides of the membrane. All states are noted as described in the text, e.g., the structural-and-functional state E_O_, whose external side is open and corresponds to S_6_ identified in the polarization study, is denoted as ^6^E_O_ (bottom middle). The left and right portions describe the transport of the intracellular ligand L_I_, e.g., Agm^+^ or a nonradioactive ligand, and the extracellular ligand L_E_, e.g., Arg^2+^ or a radioactive ligand, respectively. The middle portion described the transition of apo AdiC among eight identified states differing in conformation or energy. Transition rate constants *k*_i,j_, associated with arrows, indicate the reversible transitions between corresponding states *i* and *j* (Tables 1-4). For distinction, the transition involving the binding and unbinding of a ligand is labelled with *K_D_* = *k*_off_/*k*_on_.

### Determination of rate constants

We will use apo *S*_1_ and *S*_5_ as examples to show how we extracted rate constants among all relevant states for a given ligand condition (right side of the middle portion in Fig. 3). The above described fit of the double exponential function to the dwell-time distributions of *C*_1_ (*S*_1_+*S*_5_) yielded three independent quantities: the so-called time constants of the first (fast) and the second (slow) components (*τ*_1_ = 1/*λ*_1_ and *τ*_2_ = 1/*λ*_2_), and the fraction (*f*) of the amplitude of the first exponential component relative to the total. According to Eqs. 1–3 below (see Methods for their derivations), from *λ*_1_, *λ*_2_ and *f*, we could extract the forward and backward rate constants (*k*_15_ and *k*_51_) between *S*_1_ and *S*_5_, as well as *k*_1_ that represents the sum of the three rate constants *k*_1,j_ (*j* = 2,3,4) for the three parallel pathways of *S*_1_ to *S*_2_, *S*_3_ and *S*_4_ (Fig. 3 and Fig. S2A):

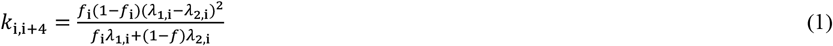

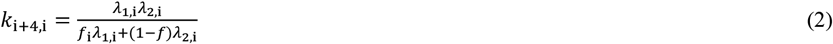

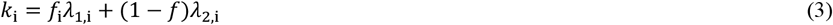

where *k*_1_ is generically denoted as *k*_i_, *k*_1,5_ as *k*_i,i+4_, and *k*_5,1_ as *k*_i+4,i_.

The determination of *k*_1,5_, *k*_5,1_ and *k*_1_ could, in principle, be obtained by the same exponential analysis of the time-dependent changes in state distributions acquired from an ensemble preparation. However, to further extract *k*_1,2_, *k*_1,3_, and *k*_1,4_ from *k*_1_ requires the information regarding the connectivity between *S*_1_ and *S*_2_, *S*_3_ or *S*_4_, namely, the observed probabilities (i.e., normalized frequencies) of transitioning from *S*_1_ to *S*_2_, *S*_3_ or *S*_4_, dubbed state-to-state transition probabilities (*p*_i,j_). It would be extremely challenging to obtain such information from an ensemble preparation, if possible. As described below, all required information to extract the three rate constants from *k*_1_ can be straightforwardly obtained from a single-molecule analysis.

For these parallel transitions, the total effective rate *k*_1_ of exiting *S*_1_ would be the sum of the rate constants *k*_1,2_, *k*_1,3_ and *k*_1,4_. Generally, the rate (*k*_i_) of exiting state *S*_i_ to one of *m* possible states is expressed as:

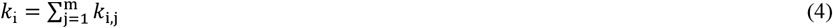

where *k*_i,j_ is given by

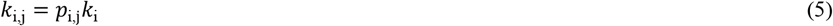

Thus, individual *k*_i,j_ could be determined from the calculated rate *k*_i_, once all relevant *p*_i,j_ values were determined. We calculated these *p*_i,j_ values from the ratio of the observed number of *C*_i_-to-*C*_j_ transitions (*n*_i,j_) and the total number of *C*_i_ events (*N*_i_):

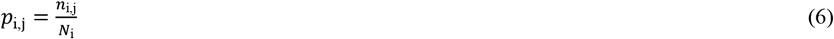

where 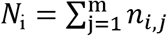. From Eqs. 5 and 6, we obtain the following equation for the direct calculation of *k*_i,j_ from *k*_i_, *n*_i,j_ and *N*_i_:

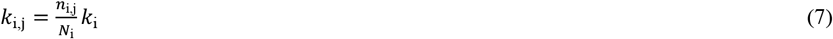

In doing so, we could determine rate constants *k*_1,2_, *k*_1,3_ and *k*_1,4_, in addition to *k*_1,5_ and *k*_5,1_ already calculated using Eqs. 1 and 2. In the same way, we also determined the remaining 15 rate constants using *τ*_1,i_, *τ*_2,i_, *f*_i_ and *p*_i,j_ of *C*_2_(*S*_2_+*S*_6_), *C*_3_ (*S*_3_+*S*_7_) and *C*_4_ (*S*_4_+*S*_8_) for the apo condition.

Furthermore, as for the apo condition, we also performed the analysis on data collected in the presence of 13 Arg^+^ concentrations or 11 Agm^2+^ concentrations. The values of *τ*_1_, *τ*_2_, *f* obtained in all examined concentration of Arg^+^ and Agm^2+^ are plotted in Fig. 4, and those of *p*_i,j_ in Fig. 5. From these four parameters for each of the four conformational states *C*_1_-*C*_4_, we calculated the 20 apparent rate constants ^app^*k*_i,j_ for each ligand condition and plotted them against the concentration of Arg^+^ or Agm^2+^ (Fig. 6). Furthermore, fitting Eq. 8 given below to all of the plots in Fig. 6, we obtained the rate constant for apo (*k*_i,j_), Arg^+^ bound or Agm^2+^ bound AdiC (^L^*k*_i,j_) all summarized in supplementary tables 1–4.

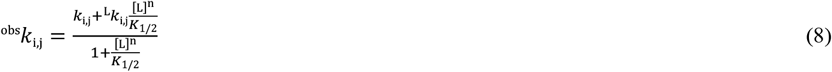

**Figure 4.**
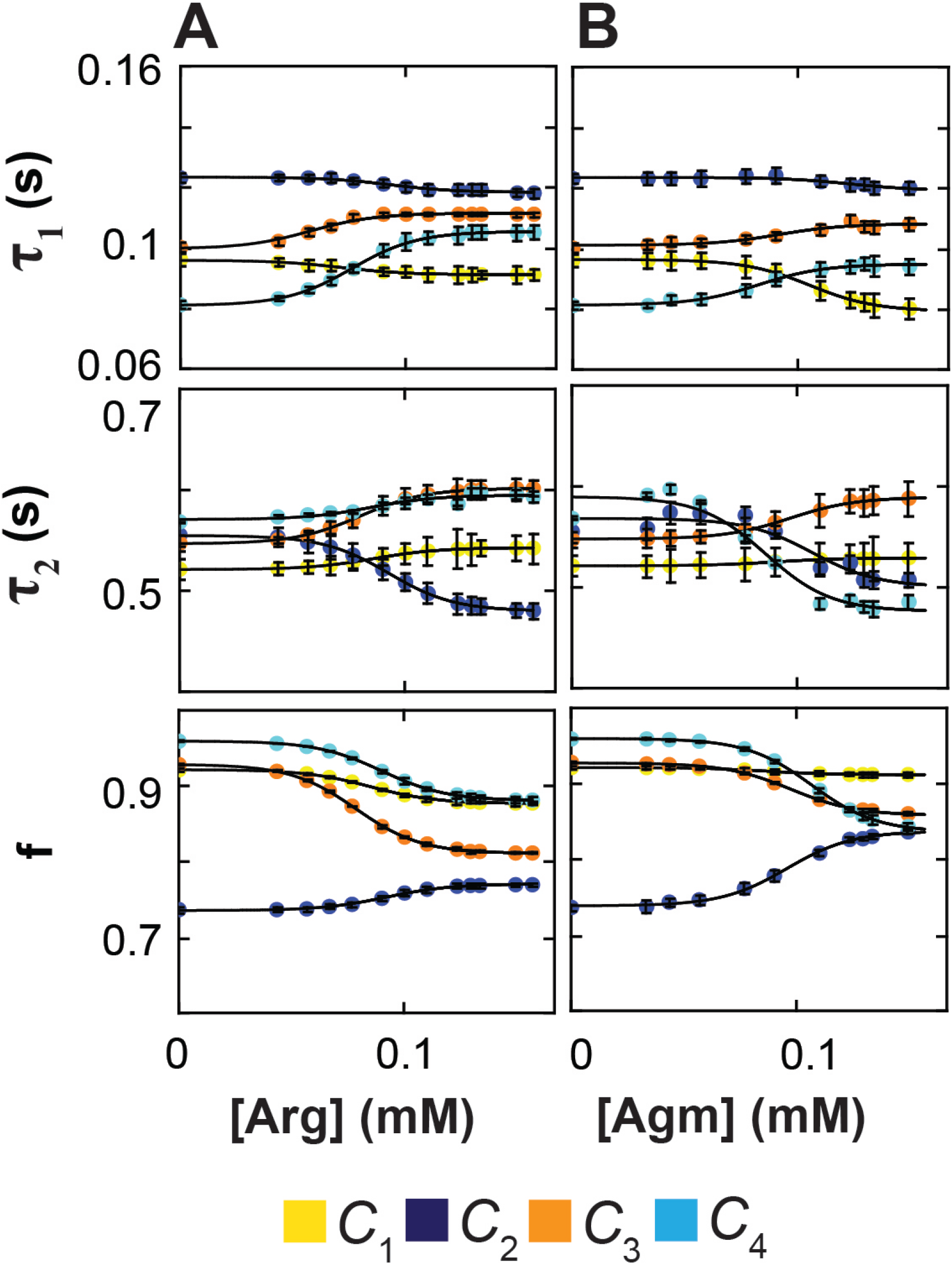
Ligand dependence of exponential fitting parameters. (A and B) Time constants *τ*_1_ and *τ*_2_ of the double exponential components (top and middle) and the relative amplitude *f* of the first component (bottom) plotted against the concentration of Arg^+^ (A) or Agm^2+^ (B). All parameters were obtained through double-exponential fits to dwell time distributions as shown in Fig. 2. The curves superimposed on the data are fits of Eq. 8. All data are plotted as mean ± sem, and symbols are color-coded for states.

**Figure 5.**
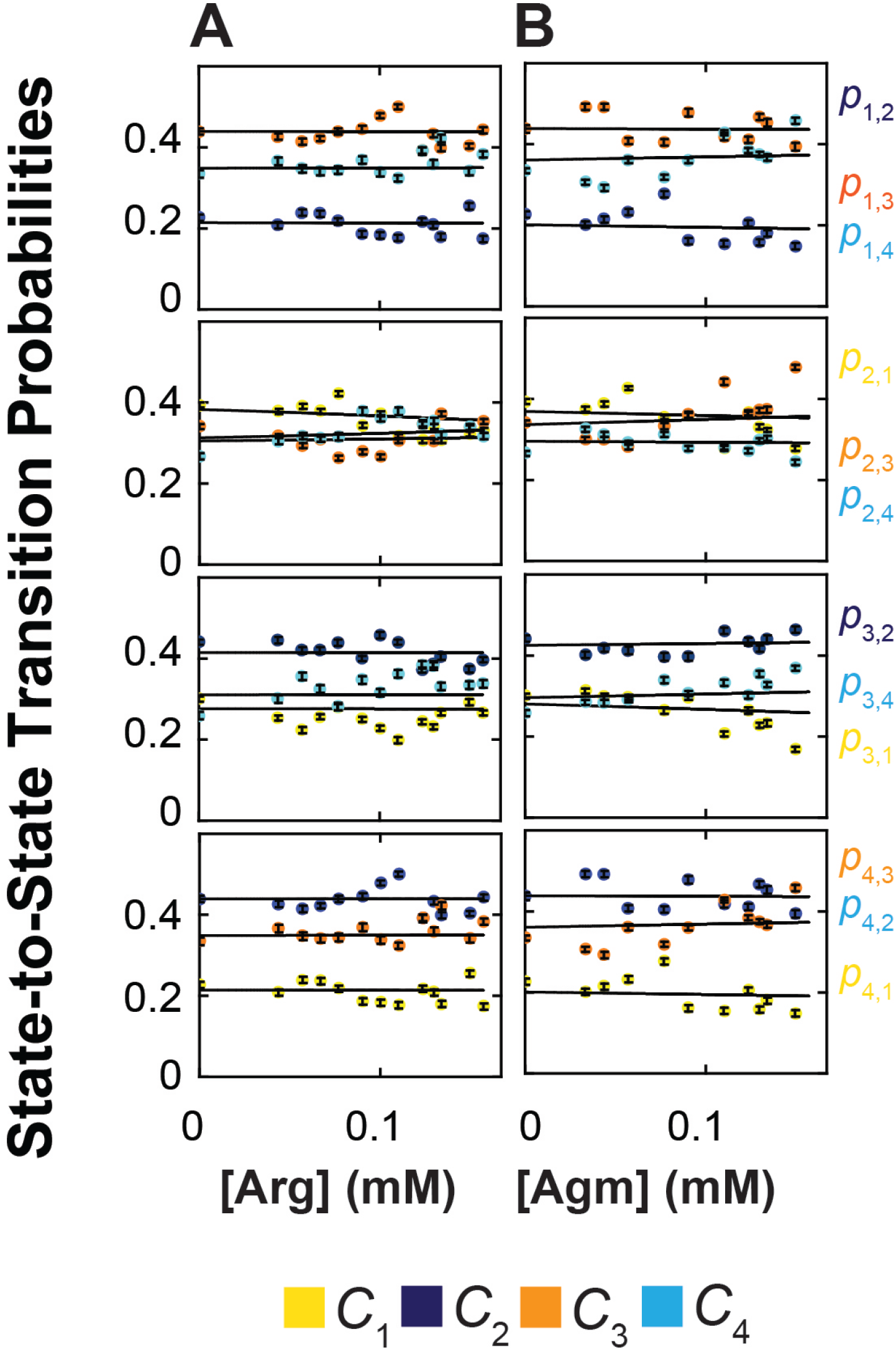
Probabilities of state-to-state transitions. (A and B) Probabilities of state-to-state transitions plotted against the concentration of Arg^+^ (A) or Agm^2+^ (B). The lines superimposed on the data correspond to the means of the respective groups of data. All data are plotted as mean ± sem, and symbols are color-coded for states.

**Figure 6.**
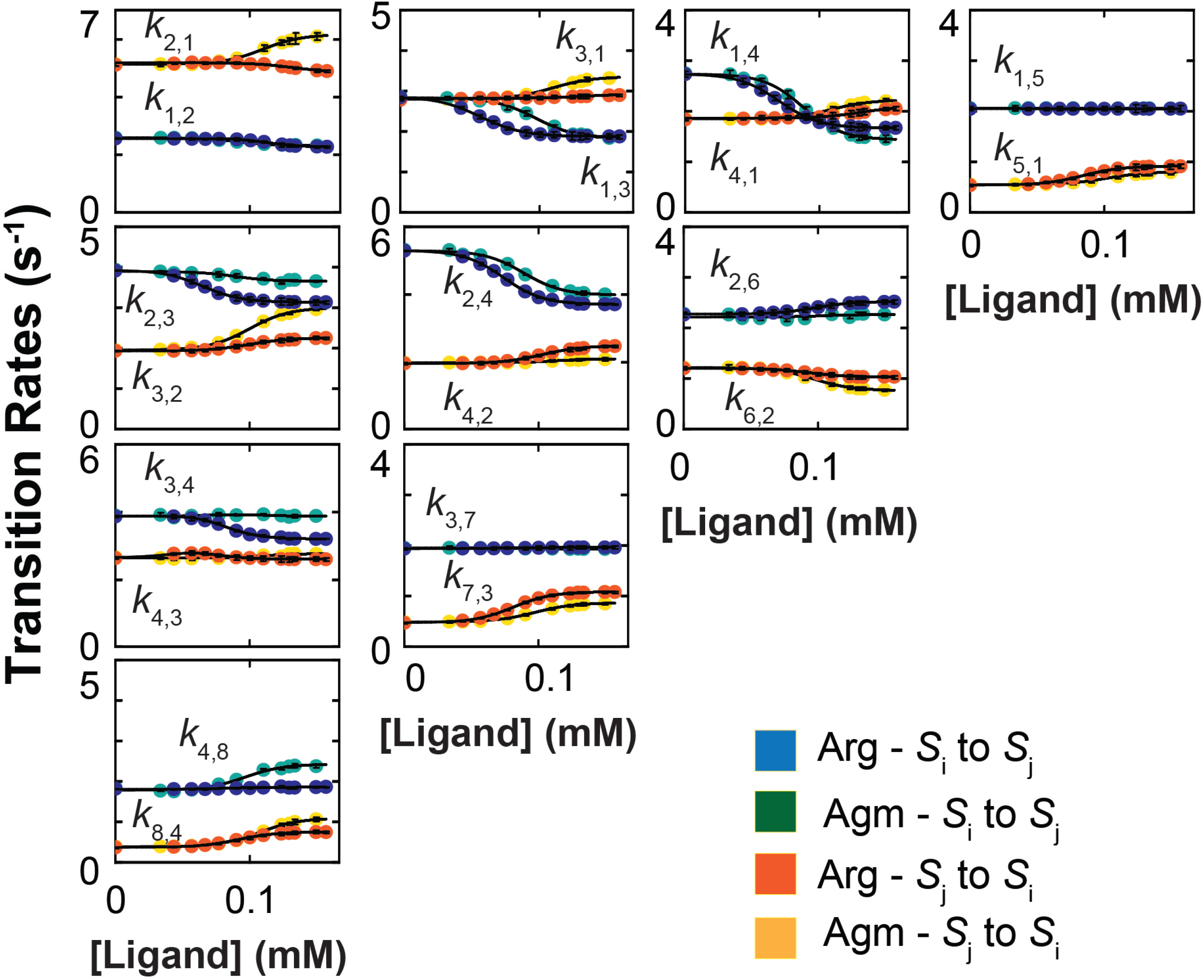
Dependence of apparent transition-rate constants on the concentration of ligands. Apparent rate constants plotted against the concentration of Arg^+^ or Agm^2+^, obtained as described in the text. The data symbols for (forward) *k*_i,j_ or (backward) *k*_j,i_ rate constants are colored blue or red in the case of Arg^+^ whereas they are colored green or orange in the case of Agm^2+^. The curves superimposed on the data are fits of Eq. 8. All fit parameter values are given in Tables 1-7. All data are plotted as mean ± sem.

The fits to the Arg^+^ and Agm^2+^ data were coupled such that a single set of rate constants for the apo condition was yielded.

To determine all K_D_ values, we first used Eq. 42 to calculate the probabilities of the eight energetic states *S*_1_-*S*_8_ (Fig. 7 and supplementary table 5) from the corresponding ^app^*k*_i,j_ values (Fig. 6) for each ligand condition. The *K_D_* values for individual states (supplementary table 6) and those of other equilibrium constants (supplementary table 7) were then determined by a global fit to the eight plots for Arg^+^ or Agm^2+^ in Fig. 7 with the equation

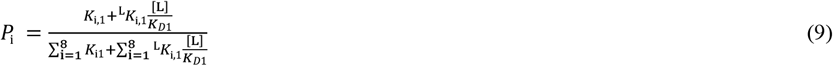

where *S*_1_ is used as a reference for other states (*S*_i_) to define equilibrium constants:

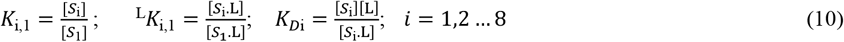

**Figure 7.**
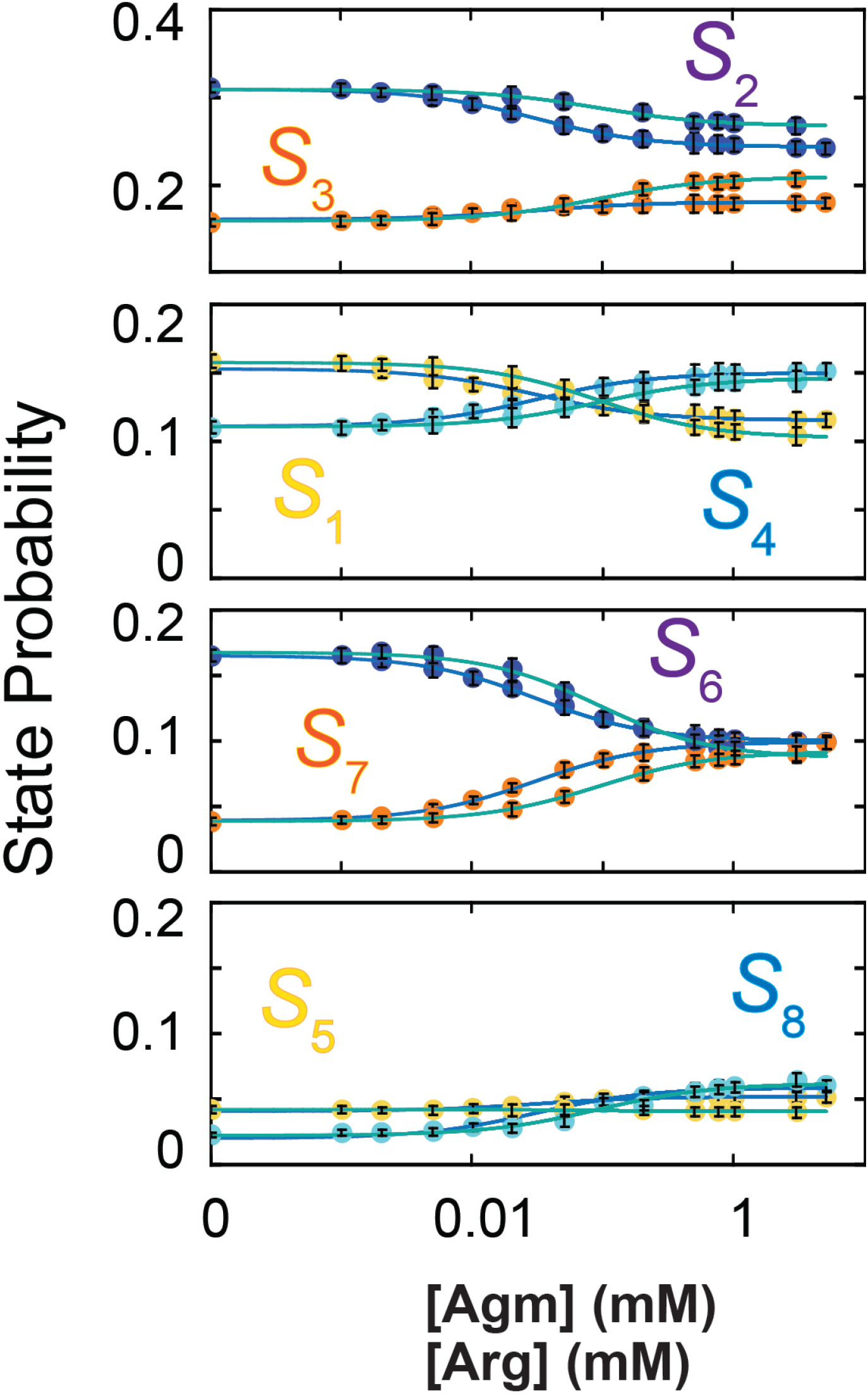
Dependence of state probabilities on the concentration of ligands. Probabilities of states *S*_1_ – *S*_8_ (yellow for *S*_1_ and *S*_5_; blue for *S*_2_ and *S*_6_; orange for *S*_3_ and *S*_7_; cyan for *S*_4_ and *S*_8_) are plotted against the concentration of Arg^+^ (with fitted curves in blue) or Agm^2+^ (with fitted curves in green). The probability values are calculated from the rate constants plotted in Fig. 6 using Eq. 42. The curves superimposed on the data are fits of Eq. 9. All fit parameter values are given in Tables 1-7. All data are plotted as mean ± sem.

The *K_D_* values of *S*_6_ and *S*_7_ that characterize the binding of external or internal Arg^+^ or Agm^2+^ are practically consequential here; they and the 60 rate constants fully determine the 24-state model of AdiC (Fig. 3).

### Characteristics of the 24-state transporter model

The Miller group has examined the substrate dependence of the rate of Arg^+^ or Agm^2+^ uptake into lipid vesicles containing purified AdiC (Tsai et al., 2012). The initial concentration of a given examined substrate type inside the vesicles was always the near saturating concentration of 5 mM whereas the bathing medium contained a varying initial concentration of radio-labeled substrate of the same type.

To simulate their uptake experiments by direct calculations, we used the solutions to a system of 24 differential equations (Eq. 44), one for each state in the 24-state model where L_E_ represents radioactive substrate and L_I_ represents non-radioactive substrate (Fig. 3). As an analytical form of the model, this system is constrained by the 60 determined rate constants for the transitions among the states in the model plus four *K_D_* values. Operationally, each *K_D_* needs to be split to the on- and off-rate constants (*k*_on_ and *k*_off_). For all cases, the substrate association- and-dissociation process is treated as a rapid equilibrium. All *k*_on_ values set to the near diffusion-limited 10^7^ M^−1^ s^−1^, and *k*_off_ values, calculated from the respective *k*_on_*K_D_*, are much greater than the other 60 rate constants; none of the calculated *k*_off_ values would have any meaningful impacts on calculated transporting rates. As shown in Fig. 8A-C, the calculated relations between the transporting rate and the substrate concentration are comparable to those experimentally observed. Consequently, the values of apparent maximum uptake rate (*k*_max_) and the concentrations (*K*_1/2_) at half of *k*_max_ from fitting the calculated and observed relations are also comparable (Table 8). Conceptually, because the calculations assume that the transport of each substrate molecule is coupled to AdiC’s conformational transitions, the comparability between the calculated and observed *k*_max_ values strongly supports the notion that AdiC is a facilitating transporter but not a channel, as the latter would allow not only one but numerous molecules to pass through its pore during each adoption of an open conformation.

**Figure 8.**
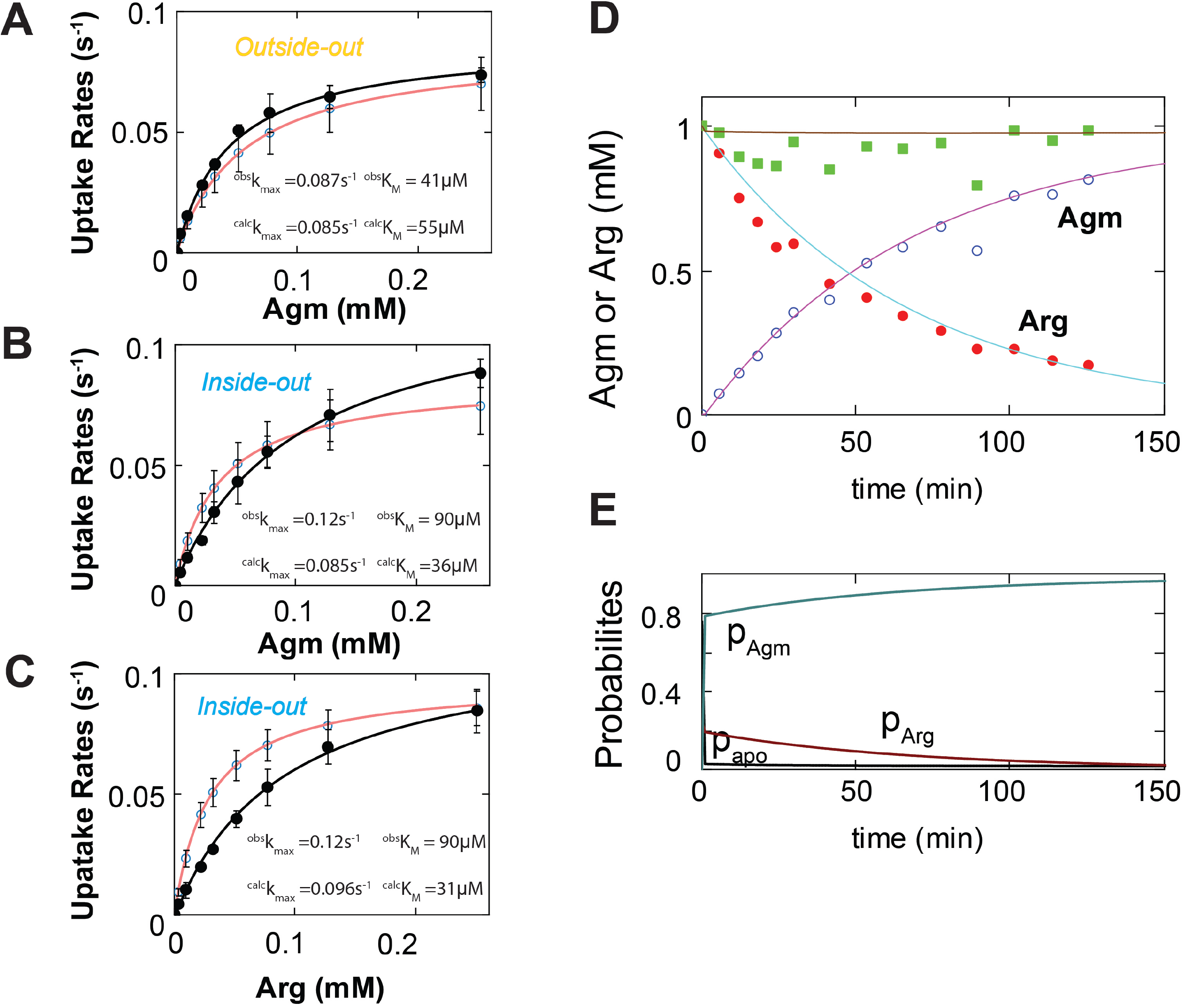
Observed and calculated rates of AdiC-mediated ligand uptakes into lipid vesicles and ligand transport across bacterial membranes. (A-C). Net ligand uptake rates plotted against the ligand concentration: Agm^2+^ by AdiC in the outside-out (A) or inside-out (B) orientation and Arg^+^ by AdiC in the outside-out orientation (C; see Fig. S1 for the comparison and comments regarding the inside-out orientation). The experimental data(Tsai et al., 2012) (mean ± sem) and their simulations (mean ± sem) were plotted as closed and open circles, respectively. The curves, which are superimposed on the symbols that represent the experimental and simulated values, correspond to the fits of the Michaelis-Menten equation. All fit parameters are presented in Table 8. D. The observed concentration of Arg^+^ (red dots), Agm^2+^ (blue open circles) or their sum (green squares) in the bacteria culture medium plotted against observation time(Iyer et al., 2002). The curves overlaid on the experimental data represent the calculated Arg^+^ (green) and Agm^2+^ (pink) concentrations; the red line corresponds to the sum of the calculated Arg^+^ and Agm^2+^ concentrations. All calculations were performed as described in Methods. **e**. The calculated probability of AdiC in apo states (black curve) or in Arg^+^- or Agm^2+^-bound states (blue or maroon curve) plotted against time, where the probability of AdiC in the apo states drop steeply within the first 1 min.

A common necessary criterion for coupled countermovement of substrates is equal amounts of substrates being transported in both directions. The experimentally observed amount of Arg^+^ uptake into the bacteria was very similar to that of Agm^2+^ coming out of bacteria and, consequently, the sum of Arg^+^ and Agm^2+^ in the medium remained approximately constant over time (Iyer et al., 2002). The calculations using our AdiC model, in which indirect transitions occur between E_o_ and I_o_ without any ligands bound and thus their countermovement is not obligatorily coupled, yield comparable relations (Fig. 8D). This is because substrates concentrations on both sides of the cell membrane became sufficiently high immediately after the start of the experiment. Consequently, the probabilities of being in the apo states should be practically negligible and the countermovement of substrates was in an apparently coupled manner (Fig. 8E).

### An integrative 4D model of AdiC

In the framework of the kinetic model, we can generate a temporal template to connect the existing experiment-based structural models of individual conformational states, and thereby create an integrated 4D transporter model for a chosen ligand condition. The template was simulated on the basis of the distributions of the experimental dwell times of individual energetic states. In a simulation of the time courses of state transitions in the transport cycle under a given ligand condition, the initial state was designated as *S*_7_ (*I*_o_). Time-dependent state-to-state transitions are governed by the corresponding rate constants organized with a 24 × 24 transition matrix **Q** (Eq. 40). AdiC stays in *S*_7_ over a simulated dwell time (*t*_sim_) before transitioning to one of the three connected states: *S*_3_, arginine-bound *S*_7_ or agmatine-bound *S*_7_ with the corresponding rates (Fig. 3). *t*_sim_ was obtained by randomly drawing from an exponential distribution of dwell times for *S*_7_, which is defined by the rates of exiting to the three states. When the accumulated time of dwelling in *S*_7_ became equal to or greater than *t*_sim_, AdiC would transition from *S*_7_ to one of the three connected states, which was determined by the outcome of a random draw from a multinomial distribution defined by the three state-to-state probabilities (Fig. 5A,B). The *t*_sim_ of dwelling in this second state and its termination point were determined as described above. These steps were repeated until the end of the entire simulation. A temporal template obtained from such a simulation is shown in Fig. 9A.

**Figure 9.**
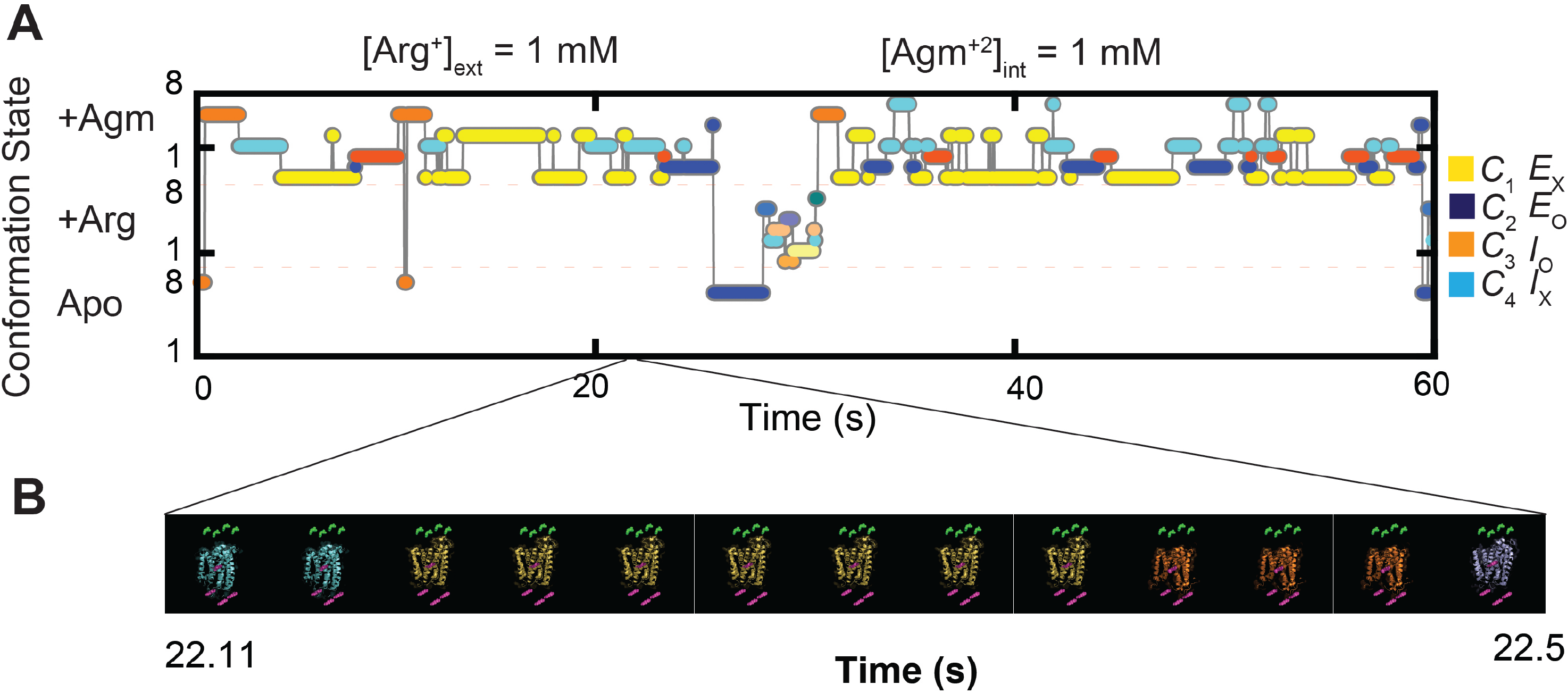
A simulated time course of conformational transitions of AdiC and a segment of a movie illustrating a 4D model of AdiC. A. Time course of AdiC transitioning among four types of conformations, simulated as described in the text for the presence of 1 mM extracellular Arg^+^ and 1 mM intracellular Agm^2+^. For clarity, AdiC’s presence in the apo, Arg^+^-bound and Agm^2+^- bound forms are indicated by individual filled circles in the lower, middle and upper sections of the panel, respectively; its adoptions of *C*_1_ (*S*_1_ + *S*_5_) are colored yellow, *C*_2_ (*S*_2_ and *S*_6_) colored blue, *C*_3_ (*S*_3_ and *S*_7_) colored orange and *C*_4_ (*S*_4_ and *S*_8_) colored cyan. B. A segment of the movie exhibiting a 4D model of AdiC transitioning among four types of conformations while transporting ligands. The movie was generated using the simulated time course shown in A as the temporal template to connect the structures of different states, as described in the text. The shown movie frames correspond to the segment of simulated time course from 2211 ms to 2250 ms.

In terms of the structural components for building a 4D model, the structures of AdiC in the E_O_ (pdb 7o82) and E_X_ (pdb 3l1l), but not I_O_ or I_X_, states have been determined. Given that AdiC shares the same type of structural fold with the BasC (pdb 6f2g) and ApcT (pdb 3gia) transporters, we used the structures of BasC or ApcT in the I_O_ or I_X_ states as temporary proxies for those of AdiC. The substrates are placed in accordance with those in AdiC.

The resulting integrative 4D model summarizes our understanding of the conformational mechanism underlying substrate transport in the spatial and temporal dimensions, which is fully defined by: ***i***. the 24-state kinetic model documented by the state diagram (Fig. 3), analytically expressed by the solutions to a system of 24 differential equations (Eq. 44) and fully defined by 60 *k*_i,j_ (supplementary tables 1–4) and 4 *K*_*D*i_ (supplementary table 5); ***ii***. the crystal structure models of the E_O_ and E_X_ states of an AdiC monomer (PDB 7o82 and 3l1l) and those of the I_O_ and I_X_ states of BasC and ApcT proper (PDB 6f2g and 3gia), respectively; ***iii***. the state-specific orientation information of helix-6A that directly links the corresponding states in the structural and kinetic models. To effectively exhibit the resulting 4D transporter model, we generated a movie to illustrate the mechanistic characteristics of the counter-transport of Arg^+^ and Agm^2+^ in spatiotemporal dimensions (supplementary video 2 for viewing in a small window). Thirteen frames of such a movie are shown in Fig. 9B.

## Discussion

The present study reveals AdiC’s complex kinetic mechanism of transitioning among eight inherent states that differ either in conformations or relative free energy. These transitions occur spontaneously, depending on thermal energy. They enable AdiC to facilitate the transmembrane movement of Arg^+^ or Agm^2+^, and thus depend also on the substrate concentrations. The concentration gradient of either Arg^+^ or Agm^2+^ or both would perturb the system’s natural distribution of states and redistribute them to facilitate a net transmembrane flux of substrates, as illustrated by Fig. 8D. Because of the existence of spontaneous conformational transitions among apo states, after transporting one substrate molecule from one side of the membrane to the other side, AdiC could cycle its open state back to the initial side without transporting another substrate molecule in the opposite direction. Such non-obligatorily coupled countermovement of substrates would ensure that substrates on a given side of the membrane could always have access to a portion of the AdiC molecules, with or without substrates on the opposite side. Nonetheless, as shown in Fig. 8D, AdiC gives rise to apparently coupled countermovement of substrates in the presence of adequately high concentrations of substrates.

In the framework of the Michaelis-Menten model, *k*_cat_ and *K*_m_ are generally used as empirical quantitative proxies for basic functional characteristics of transporters. As detailed in Methods, the maximum net flux rate *k*_max_ for the AdiC model (Fig. 3) may be expressed as

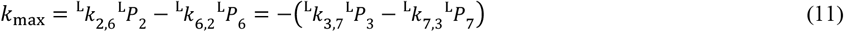

Furthermore, when the ligand concentration on one side is saturating whereas that on the opposite side is zero, *k*_max_ is at its theoretical maximum, which is the so-called *k*_cat_

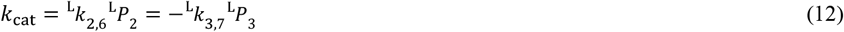

To facilitate the following discussion, *K*_m_ is expressed as

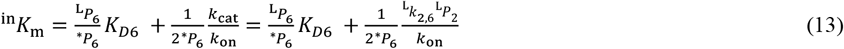

for the intracellular side, or

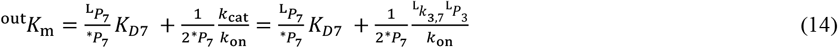

For the extracellular side; the symbol “*” in *\*P*_i_ indicates the probability of state i when the ligand concentration [L] = *K*_m_. These expressions reveal the following mechanistic information. First, according to Eq. 12, 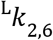 or 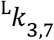 play rate-limiting roles, which are indeed among the four rate constants that are markedly smaller than the rest, under the apo, Arg^+^-bound or Agm^2+^-bound condition, seen in Fig. 10 as taller columns that represent inversed rate constants. The other two are 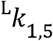 and 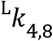 for the transitions between states not on the direct transporting paths (i.e., the *“cul-de-sac”* occluded states) and thus have minimal impacts on *k*_cat_. As expected, *k*_cat_ in theory has the same value regardless of AdiC’s orientation in the membrane (Eq. 12), so is the *k*_max_. Second, ^in^*K*_m_ and ^out^*K*_m_ for a given substrate do not need to be the same (Eqs. 13 and 14). Consistent with intuition, a change in *K_D_* alone should only affect the relevant *K*_m_ but not *k*_cat_ (Eq. 12). However, a change in *k*_cat_ would affect both ^in^*K*_m_ and ^out^*K*_m_ (Eqs. 13 and 14). Knowing the rate constants and K_D_ values in the model can predict *k*_cat_, ^in^*K*_m_ and ^out^*K*_m_; the converse is not necessarily true. Third, mechanistically defining empirical *k*_cat_ requires knowing not only the rate-limiting steps and their rate constants but also the probabilities of the relevant states (Eq. 12). The determination of these parameters in turn requires experiment-based estimates of all rate constants, an all-or-none operation that is also necessary for defining empirical ^in^*K*_m_ and ^out^*K*_m_ (Eq. 13 and 14).

**Figure 10.**
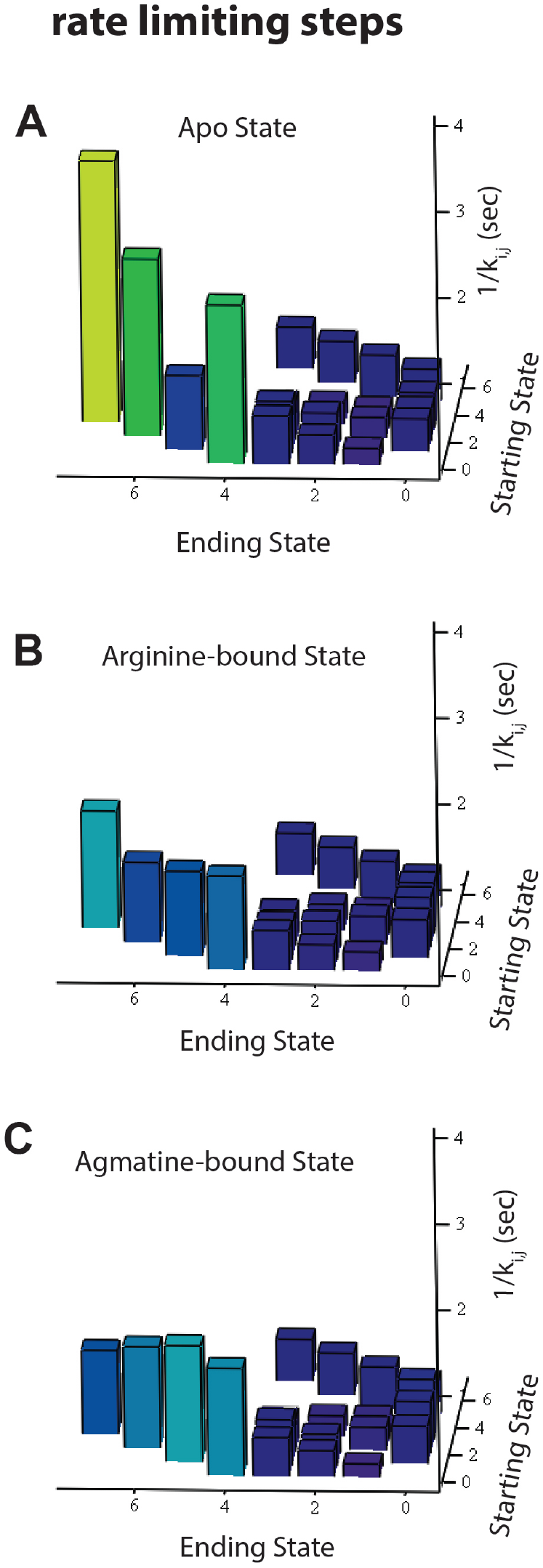
Comparison of state-to-state transition rate constants of AdiC. (A-C). The reciprocals of the rate constants (1/*k*_i,j_, z-axis) plotted against the starting state i (x-axis) and the ending state j (y-axis). The four taller columns correspond to the reciprocals of *k*_5,1_, *k*_6,2_, *k*_7,3_ and *k*_8,4_ in the absence (A) or the presence of Arg^+^ (B) or Agm^2+^ (C). The columns are color-coded according to their height, namely, changing from blue towards yellow as the height increases.

In summary, using the AdiC transporter, we have demonstrated the unprecedented capability of a high-resolution polarization-fluorescence microscopic system in solving a highly complex kinetic mechanism of transmembrane-protein-conformational dynamics by determining the 60 rate constants and 4 equilibrium dissociation constants in full. These determinations enabled us to establish an analytic kinetic model of 24 states expressed by a system of 24 differential equations that can simulate the behaviors of the transport for any specified substrate conditions. This analytic model satisfactorily predicts the previously observed transporting behaviors of AdiC. Also, the acquired temporal information is essential for understanding the mechanisms of AdiC in 4D. Combining the present temporal information and the existing structural information, we have been able to construct a fully experiment-based integrative-4D model to capture and exhibit the complex spatiotemporal mechanisms of a facilitated transport of an amino acid and its metabolite. Thus, a combination of the present method and existing structural techniques serves as an effective means to help transition structural biology, which has thus far been highly successful in the characterization of individual static structures, to an integrative form of dynamic structural biology.

## Materials and Methods

### Protein sample preparation and intensity recordings and analysis

As described in the companion manuscript, the purified recombinant protein AdiC, which had a biotin moiety linked covalently to the N-terminal biotin-ligase-recognition sequence and strep-tags in both N- and C-terminal regions, and the double G188C and S195C cysteine mutations in helix 6a, was produced using the bacterial BL-21 expression system and inserted into nanodiscs. The protein was labeled with bifunctional rhodamine (Bis-((N-Iodoacetyl)-Piperazinyl)-Sulfonerhodamine; Invitrogen B10621) via the two mutant cysteine residues and attached to streptavidin molecules adhered to the surface of a coverslip conjugated via biotinylated N-termini and strep-tags in both N- and C-terminal regions. Polarized emissions from individual bi-functional rhodamine labels, excited in the evanescent field created at the surface of the sample coverslip by a circularly polarized laser beam (532 nm), were collected via a fluorescence microscope with four polarization emission channels onto an electron-multiplying charge-coupled device camera (EMCCD), while the sample protein was immersed in a solution (pH 5) containing 100 mM NaCl, 100 mM dithiothreitol (DTT) and 50 mM acetic acid, without or with arginine or agmatine at a specific concentration.

Each intensity of the four emission components collected from a given fluorophore was a direct summation of individual pixels. I_tot_, θ, φ, and Ω were calculated using Eqs. 9–11 and 31 in the companion manuscript, respectively. Conformational transitions and states were identified in two separate steps. A changepoint algorithm (Chen and Gupta, 2001) was applied to the intensity traces to detect the transitions between conformational events, whereas a shortest-distance-based algorithm (Press et al., 2007) was used to identify the conformational states of individual events on the basis of the information regarding *θ* and *φ* angles. However, operationally, we performed the shortest-distance analysis in a cartesian coordinate system where x,y,z were calculated from the *θ* and *φ* angles along with a unity radius *r*. This analysis identified four distinct conformational state populations *C*_1_ – *C*_4_.

The experiments were performed in ten separate occasions. Data collected among these separate collections are statistically comparable and were pooled together and results in sufficiently narrow distributions as illustrated in the companion manuscript. The width of the distributions reflects both technical and biological variations. Outlier data were excluded on the following basis. First, while fluorescence intensity is expected to vary among different polarization directions, the total intensity should be essentially invariant. Therefore, any recordings whose total intensity changes beyond what were expected for random noise are excluded. Second, for a given recording, at least 12 events are required to obtain a 95% confidence level for state identification, so any traces with less than 12 events are excluded. Third, for event detection and state identification, a signal to noise ratio greater than 5 is necessary for the required minimum angle resolution. Thus, any set of intensity traces with this ratio less than 5 are excluded. The sample sizes were estimated on the basis of previous studies (Lewis and Lu, 2019c, Lewis and Lu, 2019a, Lewis and Lu, 2019b) to result in sufficiently small standard errors of mean to obtain accurate estimate of the mean. Practically, the error bars are comparable to the sizes of the symbols of data as illustrated in Figs. 4–7. The 95% confidence intervals are provided for all determined quantities in supplementary Tables 1–8. F-tests were used to evaluate single versus double exponential fits as described in a previous study (Woody et al., 2016).

### Determination of rate constants from the parameters yielded from fitting a two-exponential function to the dwell-time distributions

As the intensity signals were recorded on a camera in discrete units of time *Δt*, the dwell-time distribution for each of the four conformational states *C*_1_ – *C*_4_ was fitted with the following double exponential probability mass function (pmf)(Lewis et al., 2017):

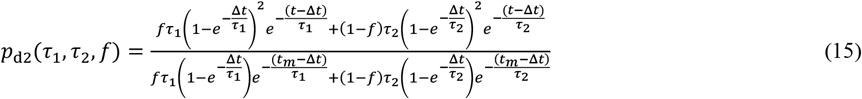

in which the missing events that are shorter than *t*_m_ were taken into consideration. Statistically, the distributions were much better fitted with such a normalized double-exponential function than a single exponential function defined as:

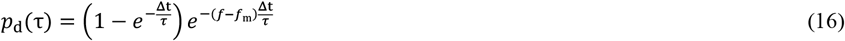

As in the Results section, we again use apo *S*_1_ and *S*_5_ as examples to derive the relations between the three fitted parameters and the three apparent underlying rate constants for a given ligand condition. The relevant part of the 24-state model in Fig. 3 is shown in Fig. S2A. In this part, *S*_1_ and *S*_5_ are reversibly connected with the rate constants *k*_15_ and *k*_51_; *S*_1_ can also transition to *S*_2_, *S*_3_ or *S*_4_ (together denoted as *S*_i_) via three parallel pathways, represented by a single step to *S*_i_ in Fig. S2A, with the combined rate constant *k*_1_, whereas the reversed three paths whose combined rate constants would be denoted by *k*_-1_ are not contributing factors of the lifetime distribution of *C*_1_ that is split to *S*_1_ and *S*_5_.

The system of rate equations for the part of the model represented in Fig. S2A is expressed as

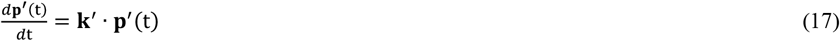

with **p**′ and **k**′ defined as:

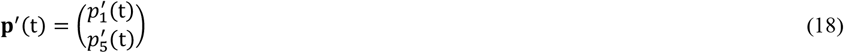

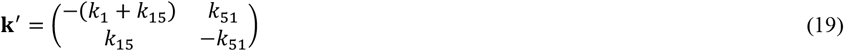

Equation 17 represents a homogeneous linear system of ordinary differential equations. The general solution for this kind of system is a set of two exponential functions expressed as:

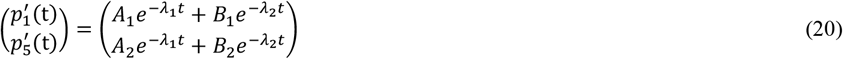

where the apparent rates *λ*_1_ and *λ*_2_ are convolutions of the rates *k*_1_, *k*_15_ and *k*_51_. The relationship between them can be obtained from the eigenvalues of **k**′ by evaluating the following determinant set to zero:

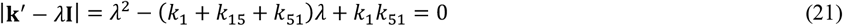

Using the quadratic equation, the two roots of *λ* are:

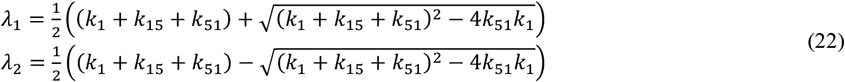

If we were observing states *S*_1_ and *S*_5_ separately, then the probabilities given by Eq. 20 would define their respective dwell time distributions. However, in the present case, *S*_1_ and *S*_5_ are observed indistinguishably in terms of spatial orientation as state *C*_1_. The expression for *C*_1_ is then given by

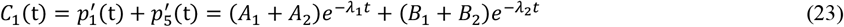

where *λ*_1_ and *λ*_2_ are defined by Eq. 22. Normalized to a probability distribution function, Eq. 23 becomes:

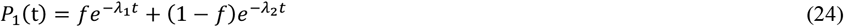

The relative amplitude of the first (fast) component, *f*, can be found from its relation to the observed average lifetime *τ*_av_ of *C*_1_ (i.e., “*S*_1_+ *S*_5_”):

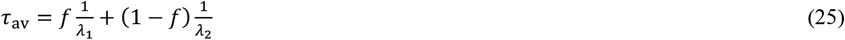

The average lifetime for the kinetic process shown in Fig. S2A can in turn be calculated from the rate constants through the following expression (Sakmann and Neher, 1995):

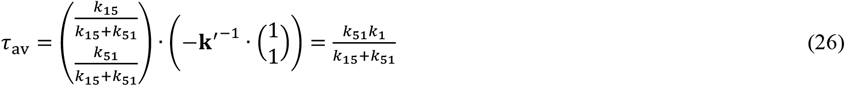

from which *f* can be calculated from the expression

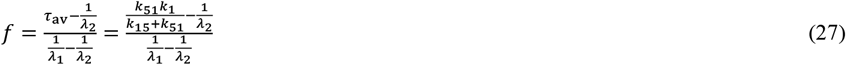

Thus, the rate constants *k*_1_, *k*_15_ and *k*_51_ can be calculated from *λ*_1_, *λ*_2_, and *f* as:

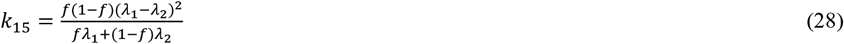

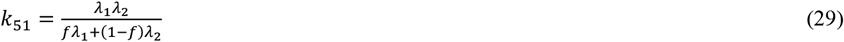

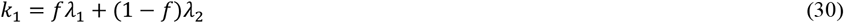

in which *λ*_1_, *λ*_2_, and *f* can be obtained from a fit of the double exponential pmf to the dwell-time distribution of *C*_1_ (“*S*_1_+ *S*_5_”). The three rate constants that sum to *k*_1_ can be further determined on the basis of state-to-state transition probabilities as described in the Results.

### Kinetic mechanism for AdiC transporter

To evaluate the substrate flux in the framework of the full model that contains eight states for apo, L_1_-bound or L_2_-bound, totaling 24 states (Fig. 3), the concurrent steady-state substrate-net-flux rates for L_1_ and L_2_ are separately calculated for a given set of conditions. When the conformational changes of AdiC are in a steady state, all sequential steps must have the same flux, defined as the difference between actual forward and backward rates, not rate constants. Thus, in the kinetic model (Fig. 3), the steady-state net flux rate of substrates of a given ligand, e.g., L_1_, on both sides of the membrane can be expressed as:

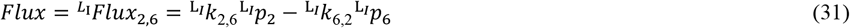

The first-order rate constants 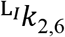 and 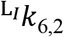 can be determined as described in the Results section. As usual, the difficulty of determining the probability of 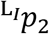 or 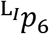 at a specified steady-state condition is underscored by the fact that each of them is defined by the solution of a differential rate equation, a solution containing all rate constants that specify the relations among all states in the model. The calculation of these probabilities is described blow.

At any given time, AdiC may exist in any one of 24 states: 8 apo, 8 L_1_-bound and 8 L_2_-bound states (Fig. 3). The probabilities of these states are represented by the column vectors ^apo^p(t), 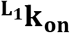 and 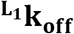:

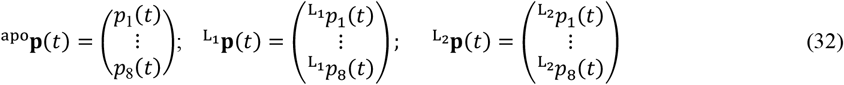

When in a system with all three types of states, the probabilities are expressed by the combined vector **p**(*t*):

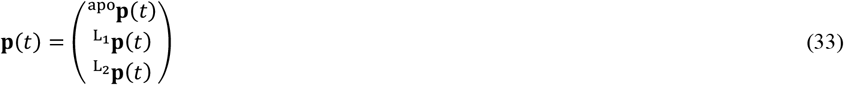

Determination of the probabilities of the 24 states will be considered in steps, with the 8 apo states alone first. The 8-state model shown in Fig. S2B, each observed conformational state (e.g., C_1_) has two kinetically connected energetic states (e.g., *S*_1_ and *S*_5_). The transition matrix for the cycle among *S*_1_ – *S*_4_ is defined as **k**:

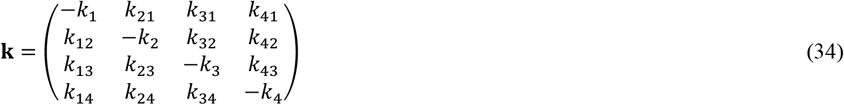

where *k*_i_ (*i* = 1…4) is as defined in the Results section, e.g., *k*_1_ = *k*_12_ + *k*_13_ + *k*_14_, and calculated in accordance with the definition given in Eq. 4, whereas the rate constants *k*_i,j_ are calculated from *k*_i_ and *p*_i,j_ using Eq. 5, in which *p*_i,j_ is calculated using Eq. 6. States 5 – 8 only have one connection each to the main cycle, i.e. *S*_1_ to *S*_5_, *S*_2_ to *S*_6_, *S*_3_ to *S*_7_ and *S*_4_ to *S*_8_ (Fig. S2B), described by one of 4 additional forward rate constants *k*_i,i+4_ and one of four reverse rates, *k*_i+4,i_ that are calculated in accordance with Eqs. 28 and 29. These rate constants can be represented in the diagonal matrices **k_ab_** and **k_ba_**:

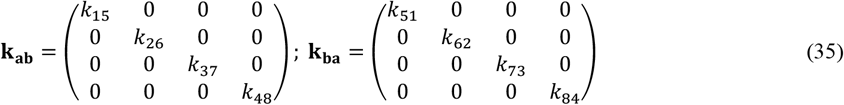

The 8X8 transition matrix ^apo^**Q** for the apo state is made up of the rates *k*_i,j_, *k*_i,i+4_ and *k*_i+4,i_:

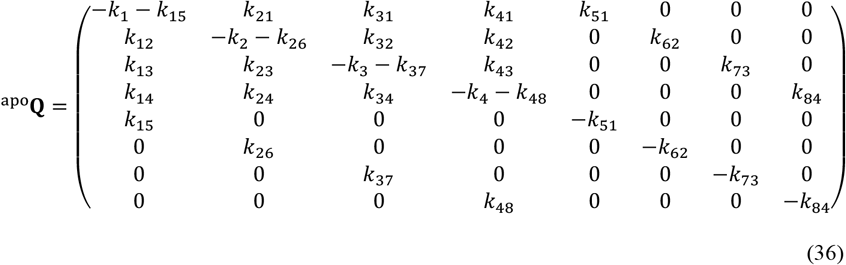

For a more concise expression, this eight state system can alternatively be represented by a system with two ensemble-state components *A* (*S*_1_ – *S*_4_) and *B* (*S*_5_ – *S*_8_) (Fig. S2C), with rates described by those in matrices **k_ab_** and **k_ba_** (Eq. 35). Using these two matrices and **k** from Eq. 34, ^apo^**Q** can be re-expressed as:

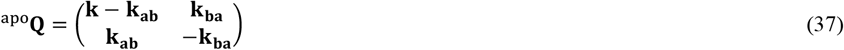

For a 16-state model including 8 apo states and 8 corresponding ligand-bound states, the transition matrices **^L1^k**, **^L1^k_ab_** and **^L1^k_ba_** would be the same forms as those used for the apo condition in Eqs. 34 and 35, with the rates replaced by ^L1^*k*_i_, ^L1^*k*_i,j_, ^L1^*k*_i,i+4_ and ^L1^*k*_i+4,i_ (Fig. 3). Entry occurs only at *S*_6_ or *S*_7_, and exit occurs at ^L1^*S*_6_ or ^L1^*S*_7_, whereas *S*_5_, *S*_8_, ^L1^*S*_5_ and ^L1^*S*_8_ are occluded “cul-de-sac” states inaccessible to solution. With these three matrices, the transition matrix for the 16 state model can be given as:

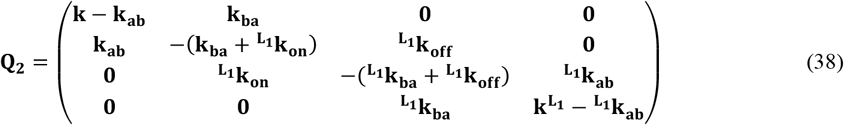

where **0** is a 4X4 matrix with zero as each element, and 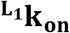 and 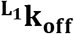 are diagonal rate matrices defined as:

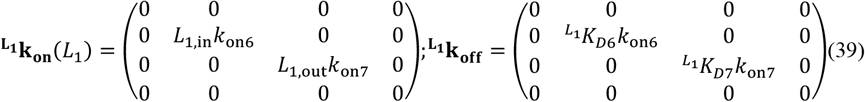

For the full 24-state model (Fig. 3) with 8 apo states, 8 L_1_-bound states and 8 L_2_-bound states, the transition matrix can be formulaically expanded to:

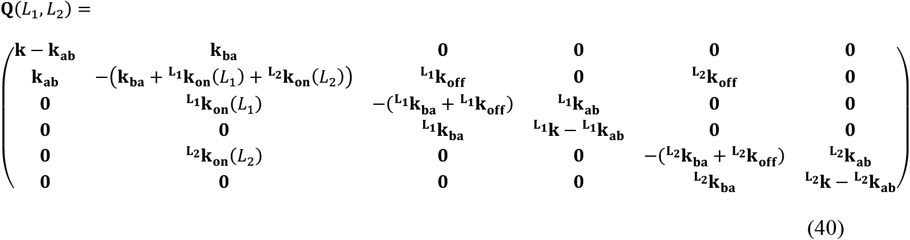

With this transition rate matrix for the 24 state model, we can simultaneously solve the linear system of 24 rate equations

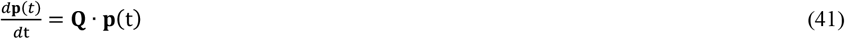

For a given steady-state condition, individual state probabilities can be calculated with Eq. 40. Mathematically, the system reaches the steady state as t → ∞, where the derivative becomes zero such that **Q** · **p**(∞) = **0**, with **0** as a zero vector. A solution to the system can be achieved by transforming the argument to the equivalent expression 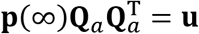, where **u** is a unit vector, **Q**_*a*_ is **Q** augmented by **u** (which would be set as an additional right-most column in **Q**_*a*_), and 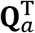 is transposed **Q**_*a*_. Solving **p**(∞) can then be done by multiplying **u** by the inverted matrix 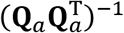, which yields(Sakmann and Neher, 1995),

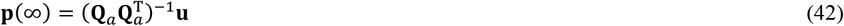

Following the determination of **p**(∞) that define the probabilities for all states including 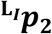 and 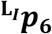, the expression for the flux 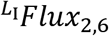 (Eq. 31) can be calculated.

### Solution of time-dependent state probabilities

The general time-dependent solution for the expression in Eq. 41 is

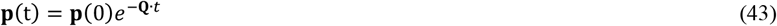

where the exponential matrix *e*^−**Q**·*t*^ can be evaluated through an expansion in accordance with the spectral theory in the form (Sakmann and Neher, 1995):

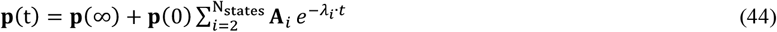

Here, **p**(∞) is from Eq. 42, *λ*_i_ are the eigenvalues of matrix **Q** (Eq. 40), determined similarly as shown in Eq. 21, and coefficient matrix **A** is calculated using the eigenvectors **X** of matrix **Q**:

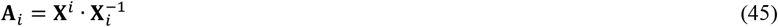

Where **X**^*i*^ is the column vector of eigenvector matrix **X**, and 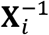 is the row vector of its inverse.

### Simulations of ligand transport for a single AdiC molecule at a steady state

The turnover rates are simulated by generating a time- and ligand-dependent flux rate:

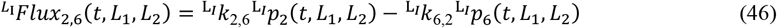

For an idealized case where the external concentration of the ligand is at a saturating level whereas the internal concentration is zero, the inward transporting rate is at the theoretical maximum, commonly called *k*_cat_. In the substrate uptake assay of the Miller group(Tsai et al., 2012), the concentration of non-radioactive substrate inside the vesicles was always 5 mM whereas the initial concentration of the radioactive substrate outside the vesicles was varied. In theory, if the concentrations of ligands on both sides of the vesicles were essentially unchanged during their assay, the uptake rate of radioactive substrate would increase to a maximum when the concentration of the radioactive substrate outside the vesicles was raised to a saturating level, which should be reflected as an increase in the initial slope of the relation between the number of transported molecules and time (Fig. S3B). This maximum rate, *k*_max_, would however be less than *k*_cat_. In some cases, the average ratio of the number of AdiC to the volumes of vesicles and their bathing medium is so high that the number of AdiC molecules is not sufficiently smaller than that of substrate molecules. In that case, the concentration gradient of the ligand drops substantially during the assay, lowering the transport rate over time. If given sufficient time, the rate would approach zero, seen as the number of transported molecules approaching a plateau with time (Fig. S3B), as is observed in their substrate uptake assays(Tsai et al., 2012).

To simulate those assays of net radioactive substrates transported into the vesicles, we calculated the transport rate with the 24-state model under the conditions that the ligand concentration was time-dependent (Fig. 8A-C and Fig. S1):

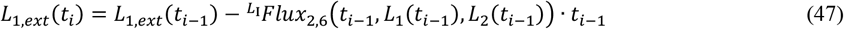

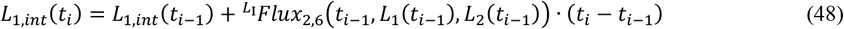

where t is a discretized series of time, and L(0) would be the starting ligand concentration. Calculation of the time dependent flux (Fig. S3B) would require iteratively calculating *L*_1,*ext*_(*t_i_*) and *L*_1,*int*_(*t_i_*), with which the flux at each time ti, i.e. 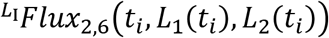 could then be calculated (Eq. 46).

We also simulated their Arg^+^-Agm^2+^ counter-transport across bacteria membranes under the conditions that the ligand concentration was time dependent(Iyer et al., 2002) (Fig. 8D). The initial Arg^+^ in the bacteria culture medium was set at 1 mM. The enzyme AdiA rapidly would convert it to an Agm^2+^ that was then transported out of the bacterium. Over time, the concentration of Arg^+^ dropped whereas that of Agm^2+^ rose with their sum staying around 1 mM in our calculation, which is in a good agreement with their experimental observation(Iyer et al., 2002) (Fig. 8D).

### Estimation of the ratio of AdiC to substrate molecules inside and outside the liposome in the previous flux assays

The Miller group performed their flux assays (Tsai et al., 2012) in 50 μL volumes, where the liposomes made up ~2% of the volume (i.e. vol_lip_ = 1 μL). Thus, the volume of solution outside the liposome (vol_out_) was 49 μL. As an example, for the case of 50 μM radioactive arginine, there should be 2.5 × 10^−9^ mol of it in vol_out_. The presence of 5 mM cold arginine inside the liposomes of total volume of about 1 μL, there should be about 5 × 10^−9^ of it in vol_lip_. The liposomes were incorporated with 6.5 μg of AdiC, translated to 6.5 × 10^−11^ mol of the dimer (100 kDa) or 1.3 × 10^−10^ mol of the monomer. In a given assay, only the molecules in either the inside-out or the outside-out orientation was examined. Thus, on a monomer basis, the effective number of AdiC was reduced by half to 6.5 × 10^−11^ mol. The estimated molar ratio of external radioactive arginine to AdiC monomer would be calculated as 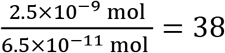, and that of the internal cold arginine as 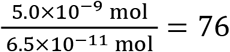. These ranges of substrate molecules per adiC are expected from Eqs. 47 and 48 to lead to a time-dependent change in the concentrations, which is manifested as a reduction in the transport rate over time (Fig. S3B). Consistent with this expectation, the number of transported molecules indeed approached a plateau with time in their assays (Tsai et al., 2012).

## Acknowledgement

This study was supported by the grant DK125521 from the National Institute of Diabetes and Digestive and Kidney Diseases.

## Data availability

A zipped source-data file for all figures is provided.

## Supplementary figure legends

**Figure S1.**
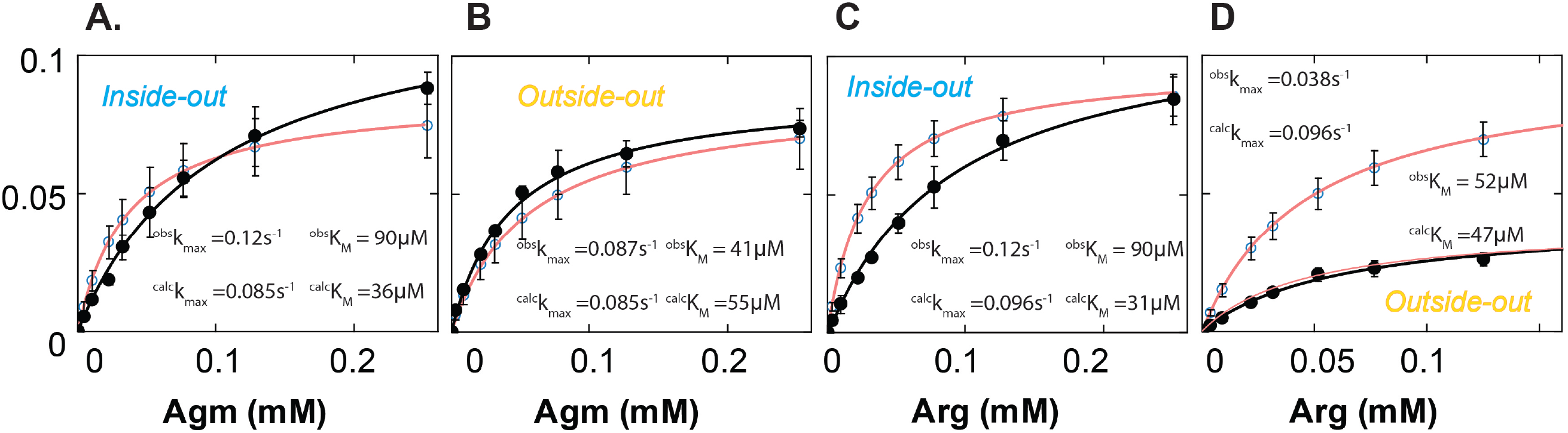
Comparison of the rates of AdiC-mediated uptakes of Arg^+^ and Agm^2+^ into lipid vesicles in different configurations. (A-D). Observed uptake rate of Arg^+^ and Agm^2+^ in the inside-out and outside-out configurations (closed circles; mean ± sem) (Tsai et al., 2012), overlaid with the simulated rates (open circles; mean ± sem). The solid curves superimposed on symbols representing the experimental and simulated values correspond to the fits of the Michaelis-Menten equation. In the presence of near saturating concentrations of a given type of substrate on both sides of the membrane, the observed maximal net-flux rates (*k*_max_) are expected to be comparable in both orientations of AdiC (see Discussion). Indeed, under three of the four conditions defined by two substrate types and two AdiC orientations, *k*_max_ values are comparable. However, *k*_max_ for Arg^+^ in the outside-out orientation is much lower, which reflects an underestimate of the maximal transporting rate or an overestimate of the effective number of AdiC molecules or both. Our calculations yield expected comparable *k*_max_ values for all four conditions. While the observed and calculated *k*_max_ values for the outside-out orientation diverge, the observed and calculated *K*_m_ are comparable: 52 versus 47 μM. This divergence is illustrated graphically with the near superposition of the observed data and the thin pink curve that represents a Michaelis-Menten equation fit to the calculated data, a curve scaled down by a factor of 2.5.

**Figure S2.**
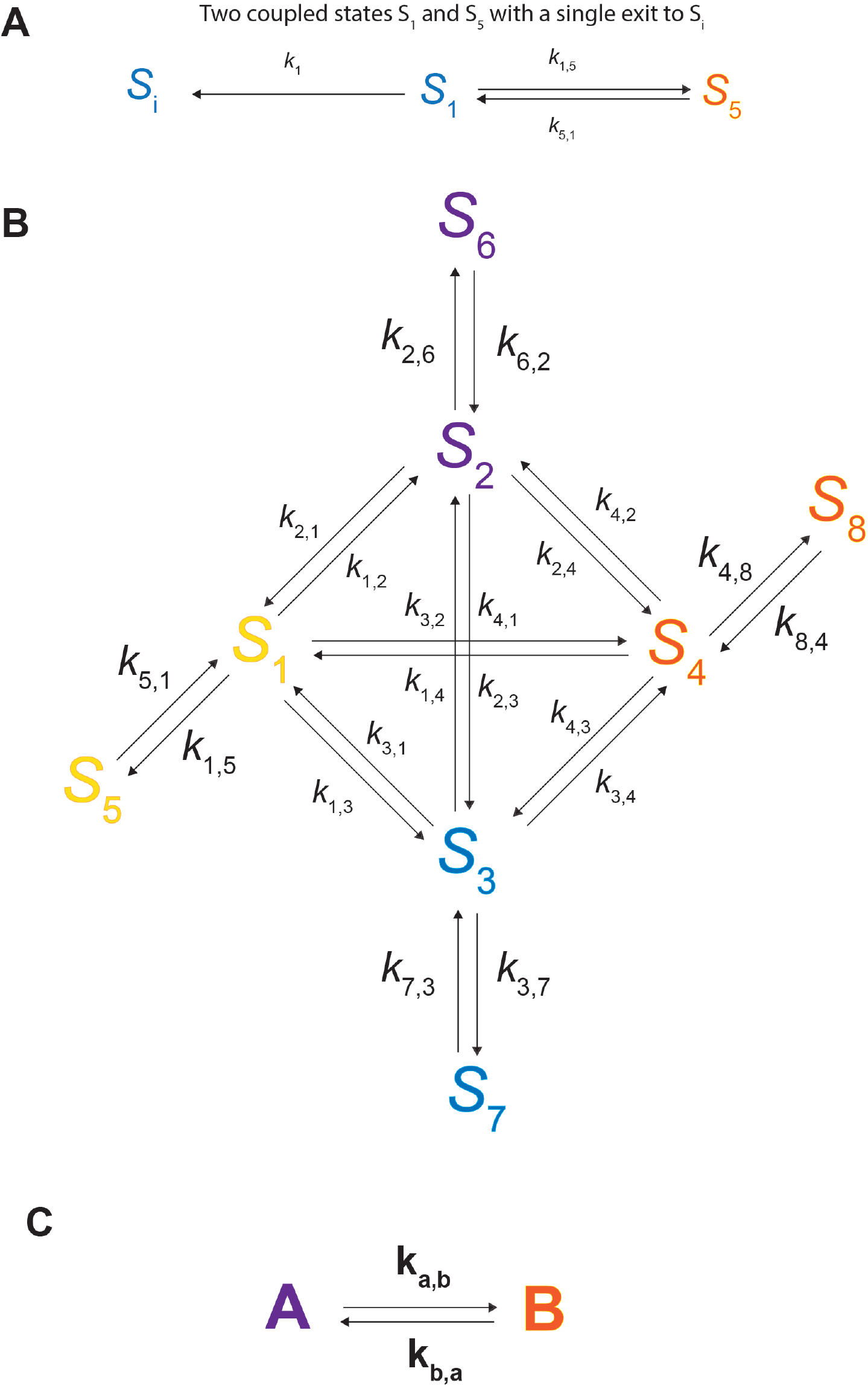
Portions of the 24-state kinetic model. (A) Diagram of the states *S*_1_ and *S*_5_ with the corresponding forward and backward rates *k*_1,5_ and *k*_5_, which are the energetic states underlying the conformation state *C*_1_. *S*_1_ transitions to an ensemble state *S*_i_ with an apparent ensemble rate constant *k*_1_. (B) Diagram of the 8 apo *S*_1_ - *S*_8_, whose transition matrix **Q**_apo_ is given in Eq. 36. **c**. The 8-state model in B is reduced to a system with two ensemble-state components, *A* and *B*, where *A* comprises the states *S*_1_ - *S*_4_ whereas *B* comprises the states *S*_5_ - *S*_8_. The corresponding transition matrix is given in Eq. 37.

**Figure S3.**
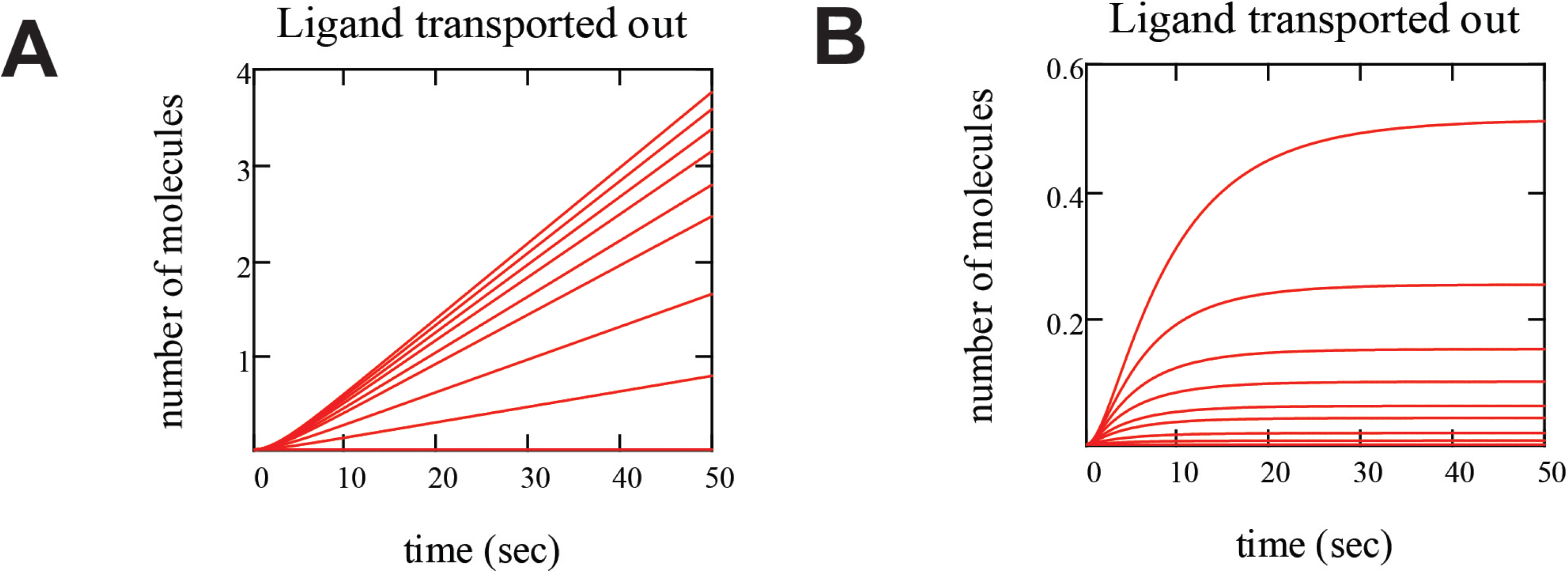
Calculated number of substrate molecules transported over time under different conditions. (A) Calculations for the condition that the molar ratio of substrate to AdiC is high such that the concentrations of substrate on the internal and external sides of the liposome remain relatively constant over time and, consequently, substrate is transported at a practically constant rate. (B) Calculations for the condition that the molar ratio of substrate to AdiC is low such that the substrate concentrations on both sides approach an equilibrium and, consequently, the transport rate drops over time and approaches zero, seen as the amount of transported substrate approaching a plateau.

## Descriptions of supplementary videos

**Video S1.** A composite video of the AdiC molecule highlighted in Fig. 1, exhibiting intensities of polarized emission of an attached fluorophore, integrated intensity values, calculated angles, and electron densities of AdiC in different conformations. ***Left***. The original recorded intensities *I*_0_, *I*_45_, *I*_90_ and *I*_135_ displayed from top to bottom. ***Middle left***. Running traces of integrated intensity values with background subtracted. ***Middle right***. Running traces of angles *θ* and *φ* along with *Ω* where each event is color-coded according to which state AdiC adopts, as determined from the state identification (Methods); *C*_1_ (yellow) corresponds to the E_X_ conformation, *C*_2_ (blue) to E_O_, *C*_3_ (orange) to I_O_ and *C*_4_ (cyan) to I_X_. ***Right***. Video of the change of the AdiC molecule among four conformational states represented by the electron densities of AdiC (PDBs 7o82 and 3l1l) and related molecules (PDBs 6f2g and 3gia), as described in the results section. At a given time point, the electron density for the state identified by the polarization study is displayed. Due to the limit of allowable file size, the video was made to be viewed in a small window of such software as Windows Media Player.

**Video S2.** Video of simulation of a counter-transport of Arg^+^ and Agm^2+^ by AdiC. As described in the results section, at a given time point, the crystal structure model for a specific state is displayed according to the template shown in Fig. 9A. The *C*_1_ (*S*_1_ plus *S*_5_) state corresponds to the E_X_ conformation, *C*_2_ (*S*_2_ plus *S*_6_) to E_O_, *C*_3_ (*S*_3_ plus *S*_7_) to I_O_ and *C*_4_ (*S*_4_ plus *S*_8_) to I_X_. Arg^+^ (shown in green) is transported from the external (top) side to the intracellular (bottom) side whereas Agm^2+^ (shown in maroon) is transported in the opposite direction. Throughout the video, external Arg^+^ and internal Agm^2+^ are both kept at constant 1 mM. The number of transported substrate molecules is assumed to be negligible with respect to the total number of available substrate molecules. Due to the limit of allowable file size, the video was made to be viewed in a small window of such software as Windows Media Player.

**Supplementary Table 1:** Lifetimes and transition probabilities for AdiC state 1 in the presence or absence of Arg or Agm

| Lifetimes (s)<sup>a</sup> |  | Transition Probabilities<sup>b</sup> |  | Transition Rates (s<sup>-1</sup>)<sup>c</sup> |  |
| --- | --- | --- | --- | --- | --- |
| $\tau_1$ | 0.096 +0.003/-0.004 | $P_{1,2}$ | 0.525 +0.036/-0.021 | $k_{1,2}$ | 5.08 +0.448/-0.274 |
| $\tau_2$ | 0.527 +0.015/-0.014 | $P_{1,3}$ | 0.286 +0.014/-0.024 | $k_{1,3}$ | 2.76 +0.205/-0.253 |
| | | | | $k_{1,4}$ | 1.82 +0.236/-0.195 |
| $f$ | 0.918 +0.001/-0.002 | $P_{1,4}$ | 0.189 +0.022/-0.019 | $k_{1,5}$ | 0.557 +0.022/-0.013 |
| | | | | $k_{5,1}$ | 2.03 +0.051/-0.057 |
| $^{\text{arg}}\tau_1$ | 0.094 +0.001/-0.0002 | $^{\text{arg}}P_{1,2}$ | 0.492 +0.035/-0.034 | $^{\text{arg}}k_{1,2}$ | 4.68 +0.354/-0.342 |
| $^{\text{arg}}\tau_2$ | 0.545 +0.116/-0.087 | $^{\text{arg}}P_{1,3}$ | 0.296 +0.029/-0.025 | $^{\text{arg}}k_{1,3}$ | 2.82 +0.285/-0.247 |
| | | | | $^{\text{arg}}k_{1,4}$ | 2.02 +0.170/-0.187 |
| $^{\text{arg}}f$ | 0.870 +0.002/-0.003 | $^{\text{arg}}P_{1,4}$ | 0.212 +0.017/-0.019 | $^{\text{arg}}k_{1,5}$ | 0.948 +0.013/-0.011 |
| | | | | $^{\text{arg}}k_{5,1}$ | 1.98 +0.055/-0.024 |
| $^{\text{agm}}\tau_1$ | 0.079 +0.005/-0.005 | $^{\text{agm}}P_{1,2}$ | 0.527 +0.019/-0.029 | $^{\text{agm}}k_{1,2}$ | 6.16 +0.354/-0.342 |
| $^{\text{agm}}\tau_2$ | 0.524 +0.026/-0.021 | $^{\text{agm}}P_{1,3}$ | 0.282 +0.027/-0.020 | $^{\text{agm}}k_{1,3}$ | 3.29 +0.410/-0.320 |
| | | | | $^{\text{agm}}k_{1,4}$ | 2.23 +0.342/-0.238 |
| $^{\text{agm}}f$ | 0.911 +0.003/-0.003 | $^{\text{agm}}P_{1,4}$ | 0.191 +0.025/-0.016 | $^{\text{agm}}k_{1,5}$ | 0.800 +0.039/-0.030 |
| | | | | $^{\text{agm}}k_{5,1}$ | 2.06 +0.082/-0.091 |

<sup>a</sup> $\tau_1$ , $\tau_2$ and $f$ are the lifetimes and amplitude obtained from a fit of Eq. 8 to the plots in Fig. 3a specific to state 1. Maximum Likelihood fits were based on a discrete two exponential pmf (Eq. 15), and performed simultaneously on state 1 dwell time distributions obtained over all ligand conditions coupled with Eq. 8. To determine confidence intervals, the data sets were re-sampled 100 times by bootstrapping, the results of which were fit as before. The upper and lower boundaries (+/-) that contain 95% of the fit values define the confidence intervals.
<sup>b</sup> $P_{1,j}$ is the probability of state 1 transitioning to state $j$ (Eq. 6). The Maximum Likelihood fit is based on a binomial pdf, with the probability defined by an equation in the same form as Eq. 8. 95% confidence interval values were calculated as in <sup>a</sup>.
<sup>c</sup> $k_{1,j}$ is the rate of state 1 transitioning to state $j$ , calculated from Eq. 5 with $k_1$ (Eq.3) and $p_{1,j}$ . 95% confidence interval values were calculated as in <sup>a</sup>.

**Supplementary Table 2:**
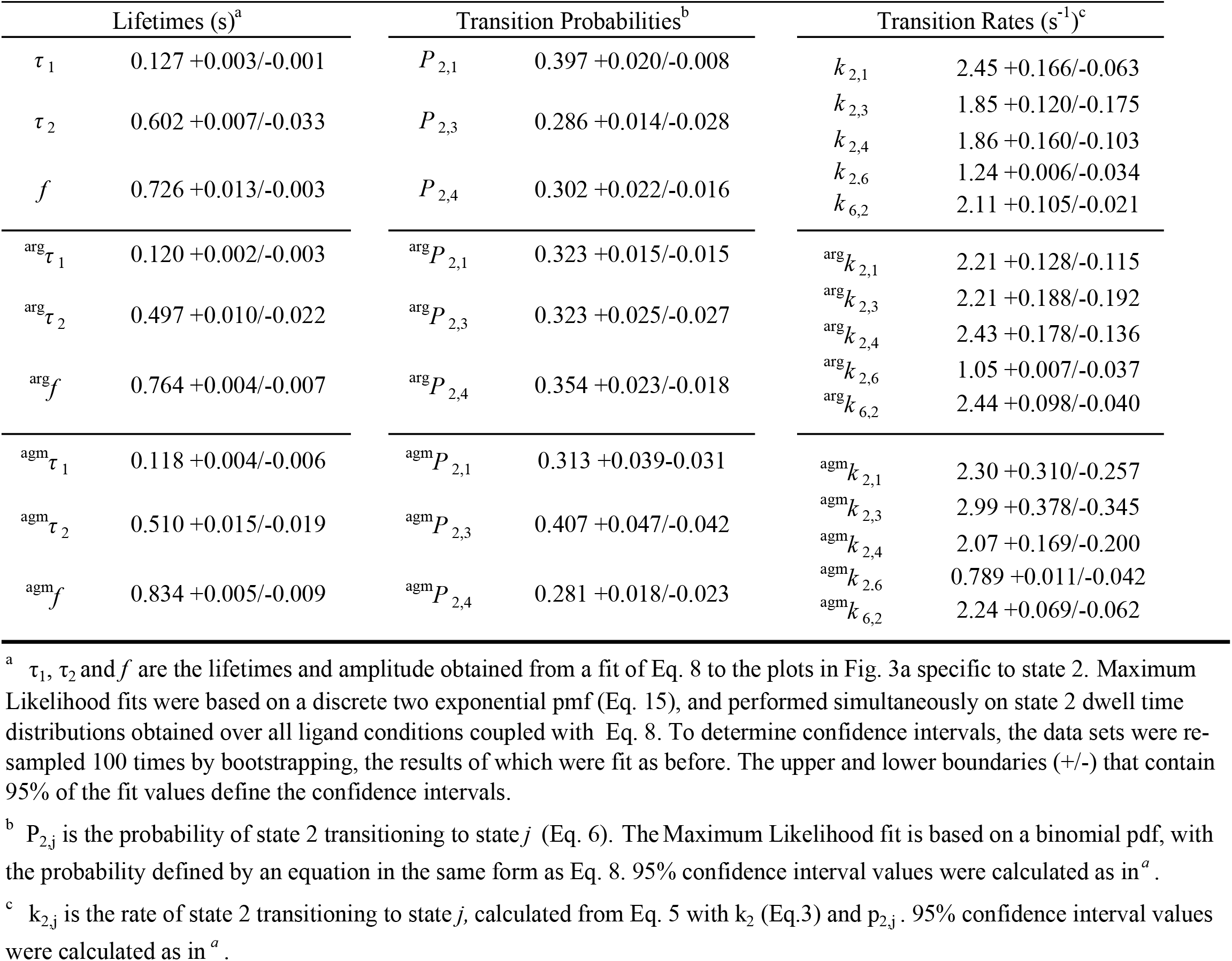
Lifetimes and transition probabilities for AdiC state 2 in the presence or absence of Arg or Agm

| Lifetimes (s)<sup>a</sup> |  | Transition Probabilities<sup>b</sup> |  | Transition Rates (s<sup>-1</sup>)<sup>c</sup> |  |
| --- | --- | --- | --- | --- | --- |
| $\tau_1$ | 0.127 +0.003/-0.001 | $P_{2,1}$ | 0.397 +0.020/-0.008 | $k_{2,1}$ | 2.45 +0.166/-0.063 |
| $\tau_2$ | 0.602 +0.007/-0.033 | $P_{2,3}$ | 0.286 +0.014/-0.028 | $k_{2,3}$ | 1.85 +0.120/-0.175 |
| $f$ | 0.726 +0.013/-0.003 | $P_{2,4}$ | 0.302 +0.022/-0.016 | $k_{2,4}$ | 1.86 +0.160/-0.103 |
| | | | | $k_{2,6}$ | 1.24 +0.006/-0.034 |
| | | | | $k_{6,2}$ | 2.11 +0.105/-0.021 |
| $^{\text{arg}}\tau_1$ | 0.120 +0.002/-0.003 | $^{\text{arg}}P_{2,1}$ | 0.323 +0.015/-0.015 | $^{\text{arg}}k_{2,1}$ | 2.21 +0.128/-0.115 |
| $^{\text{arg}}\tau_2$ | 0.497 +0.010/-0.022 | $^{\text{arg}}P_{2,3}$ | 0.323 +0.025/-0.027 | $^{\text{arg}}k_{2,3}$ | 2.21 +0.188/-0.192 |
| $^{\text{arg}}f$ | 0.764 +0.004/-0.007 | $^{\text{arg}}P_{2,4}$ | 0.354 +0.023/-0.018 | $^{\text{arg}}k_{2,4}$ | 2.43 +0.178/-0.136 |
| | | | | $^{\text{arg}}k_{2,6}$ | 1.05 +0.007/-0.037 |
| | | | | $^{\text{arg}}k_{6,2}$ | 2.44 +0.098/-0.040 |
| $^{\text{agm}}\tau_1$ | 0.118 +0.004/-0.006 | $^{\text{agm}}P_{2,1}$ | 0.313 +0.039/-0.031 | $^{\text{agm}}k_{2,1}$ | 2.30 +0.310/-0.257 |
| $^{\text{agm}}\tau_2$ | 0.510 +0.015/-0.019 | $^{\text{agm}}P_{2,3}$ | 0.407 +0.047/-0.042 | $^{\text{agm}}k_{2,3}$ | 2.99 +0.378/-0.345 |
| $^{\text{agm}}f$ | 0.834 +0.005/-0.009 | $^{\text{agm}}P_{2,4}$ | 0.281 +0.018/-0.023 | $^{\text{agm}}k_{2,4}$ | 2.07 +0.169/-0.200 |
| | | | | $^{\text{agm}}k_{2,6}$ | 0.789 +0.011/-0.042 |
| | | | | $^{\text{agm}}k_{6,2}$ | 2.24 +0.069/-0.062 |

<sup>a</sup> $\tau_1$ , $\tau_2$ and $f$ are the lifetimes and amplitude obtained from a fit of Eq. 8 to the plots in Fig. 3a specific to state 2. Maximum Likelihood fits were based on a discrete two exponential pmf (Eq. 15), and performed simultaneously on state 2 dwell time distributions obtained over all ligand conditions coupled with Eq. 8. To determine confidence intervals, the data sets were re-sampled 100 times by bootstrapping, the results of which were fit as before. The upper and lower boundaries (+/-) that contain 95% of the fit values define the confidence intervals.
<sup>b</sup> $P_{2,j}$ is the probability of state 2 transitioning to state $j$ (Eq. 6). The Maximum Likelihood fit is based on a binomial pdf, with the probability defined by an equation in the same form as Eq. 8. 95% confidence interval values were calculated as in <sup>a</sup>.
<sup>c</sup> $k_{2,j}$ is the rate of state 2 transitioning to state $j$ , calculated from Eq. 5 with $k_2$ (Eq.3) and $p_{2,j}$ . 95% confidence interval values were calculated as in <sup>a</sup>.

**Supplementary Table 3:** Lifetimes and transition probabilities for AdiC state 3 in the presence or absence of Arg or Agm

| Lifetimes (s)<sup>a</sup> |  | Transition Probabilities<sup>b</sup> |  | Transition Rates (s<sup>-1</sup>)<sup>c</sup> |  |
| --- | --- | --- | --- | --- | --- |
| $\tau_1$ | 0.101 +0.002/-0.003 | $P_{3,1}$ | 0.303 +0.025/-0.007 | $k_{3,1}$ | 2.81 +0.256/-0.092 |
| $\tau_2$ | 0.550 +0.024/-0.011 | $P_{3,2}$ | 0.417 +0.013/-0.024 | $k_{3,2}$ | 3.87 +0.190/-0.240 |
| $f$ | 0.924 +0.001/-0.001 | $P_{3,4}$ | 0.280 +0.026/-0.025 | $k_{3,4}$ | 2.60 +0.261/-0.240 |
| | | | | $k_{3,7}$ | 0.499 +0.016/-0.006 |
| | | | | $k_{7,3}$ | 1.94 +0.042/-0.081 |
| $\text{arg}\tau_1$ | 0.113 +0.001/-0.002 | $\text{arg}P_{3,1}$ | 0.247 +0.014/-0.015 | $\text{arg}k_{3,1}$ | 1.82 +0.112/-0.113 |
| $\text{arg}\tau_2$ | 0.630 +0.013/-0.018 | $\text{arg}P_{3,2}$ | 0.412 +0.018/-0.011 | $\text{arg}k_{3,2}$ | 3.04 +0.151/-0.089 |
| $\text{arg}f$ | 0.797 +0.002/-0.004 | $\text{arg}P_{3,4}$ | 0.341 +0.011/-0.018 | $\text{arg}k_{3,4}$ | 2.51 +0.101/-0.136 |
| | | | | $\text{arg}k_{3,7}$ | 1.15 +0.022/-0.019 |
| | | | | $\text{arg}k_{7,3}$ | 1.90 +0.052/-0.040 |
| $\text{agm}\tau_1$ | 0.109 +0.003/-0.005 | $\text{agm}P_{3,1}$ | 0.220 +0.037/-0.034 | $\text{agm}k_{3,1}$ | 1.77 +0.314/-0.280 |
| $\text{agm}\tau_2$ | 0.597 +0.019/-0.023 | $\text{agm}P_{3,2}$ | 0.445 +0.016/-0.034 | $\text{agm}k_{3,2}$ | 3.58 +0.236/-0.295 |
| $\text{agm}f$ | 0.866 +0.002/-0.005 | $\text{agm}P_{3,4}$ | 0.335 +0.030/-0.021 | $\text{agm}k_{3,4}$ | 2.70 +0.284/-0.188 |
| | | | | $\text{agm}k_{3,7}$ | 0.851 +0.025/-0.021 |
| | | | | $\text{agm}k_{7,3}$ | 1.89 +0.066/-0.059 |

<sup>a</sup> $\tau_1$ , $\tau_2$ and $f$ are the lifetimes and amplitude obtained from a fit of Eq. 8 to the plots in Fig. 3a specific to state 3. Maximum Likelihood fits were based on a discrete two exponential pmf (Eq. 15), and performed simultaneously on state 3 dwell time distributions obtained over all ligand conditions coupled with Eq. 8. To determine confidence intervals, the data sets were re-sampled 100 times by bootstrapping, the results of which were fit as before. The upper and lower boundaries (+/-) that contain 95% of the fit values define the confidence intervals.
<sup>b</sup> $P_{3,j}$ is the probability of state 3 transitioning to state $j$ (Eq. 6). The Maximum Likelihood fit is based on a binomial pdf, with the probability defined by an equation in the same form as Eq. 8. 95% confidence interval values were calculated as in <sup>a</sup>.
<sup>c</sup> $k_{3,j}$ is the rate of state 3 transitioning to state $j$ , calculated from Eq. 5 with $k_3$ (Eq.3) and $p_{3,j}$ . 95% confidence interval values were calculated as in <sup>a</sup>.

**Supplementary Table 4:** Lifetimes and transition probabilities for AdiC state 4 in the presence or absence of Arg or Agm

| Lifetimes (s)<sup>a</sup> |  | Transition Probabilities<sup>b</sup> |  | Transition Rates (s<sup>-1</sup>)<sup>c</sup> |  |
| --- | --- | --- | --- | --- | --- |
| $\tau_1$ | 0.080 +0.003/-0.001 | $P_{4,1}$ | 0.230 +0.023/-0.019 | $k_{4,1}$ | 2.78 +0.281/-0.260 |
| $\tau_2$ | 0.585 +0.001/-0.007 | $P_{4,2}$ | 0.445 +0.038/-0.026 | $k_{4,2}$ | 5.39 +0.466/-0.393 |
| | | | | $k_{4,3}$ | 3.93 +0.213/-0.327 |
| $f$ | 0.961 +0.002/-0.001 | $P_{4,3}$ | 0.325 +0.017/-0.023 | $k_{4,8}$ | 0.363 +0.006/-0.04 |
| | | | | $k_{8,4}$ | 1.76 +0.023/-0.001 |
| $\text{arg}\tau_1$ | 0.113 +0.001/-0.008 | $\text{arg}P_{4,1}$ | 0.195 +0.023/-0.018 | $\text{arg}k_{4,1}$ | 1.54 +0.225/-0.145 |
| $\text{arg}\tau_2$ | 0.610 +0.003/-0.016 | $\text{arg}P_{4,2}$ | 0.432 +0.029/-0.024 | $\text{arg}k_{4,2}$ | 3.41 +0.371/-0.200 |
| | | | | $\text{arg}k_{4,3}$ | 2.95 +0.320/-0.205 |
| $\text{arg}f$ | 0.870 +0.004/-0.010 | $\text{arg}P_{4,3}$ | 0.373 +0.025/-0.025 | $\text{arg}k_{4,8}$ | 0.742 +0.019/-0.003 |
| | | | | $\text{arg}k_{8,4}$ | 1.83 +0.042/-0.014 |
| $\text{agm}\tau_1$ | 0.096 +0.003/-0.005 | $\text{agm}P_{4,1}$ | 0.154 +0.028/-0.022 | $\text{agm}k_{4,1}$ | 1.38 +0.270/-0.205 |
| $\text{agm}\tau_2$ | 0.486 +0.012/-0.017 | $\text{agm}P_{4,2}$ | 0.428 +0.037/-0.033 | $\text{agm}k_{4,2}$ | 3.85 +0.430/-0.334 |
| | | | | $\text{agm}k_{4,3}$ | 3.75 +0.462/-0.365 |
| $\text{agm}f$ | 0.834 +0.006/-0.012 | $\text{agm}P_{4,3}$ | 0.417 +0.042/-0.037 | $\text{agm}k_{4,8}$ | 1.061 +0.027/-0.023 |
| | | | | $\text{agm}k_{8,4}$ | 2.37 +0.071/-0.051 |

<sup>a</sup> $\tau_1$ , $\tau_2$ and $f$ are the lifetimes and amplitude obtained from a fit of Eq. 8 to the plots in Fig. 3a specific to state 4. Maximum Likelihood fits were based on a discrete two exponential pmf (Eq. 15), and performed simultaneously on state 4 dwell time distributions obtained over all ligand conditions coupled with Eq. 8. To determine confidence intervals, the data sets were re-sampled 100 times by bootstrapping, the results of which were fit as before. The upper and lower boundaries (+/-) that contain 95% of the fit values define the confidence intervals.
<sup>b</sup> $P_{4,j}$ is the probability of state 4 transitioning to state $j$ (Eq. 6). The Maximum Likelihood fit is based on a binomial pdf, with the probability defined by an equation in the same form as Eq. 8. 95% confidence interval values were calculated as in <sup>a</sup>.
<sup>c</sup> $k_{4,j}$ is the rate of state 4 transitioning to state $j$ , calculated from Eq. 5 with $k_4$ (Eq.3) and $p_{4,j}$ . 95% confidence interval values were calculated as in <sup>a</sup>.

**Supplementary Table 5:**
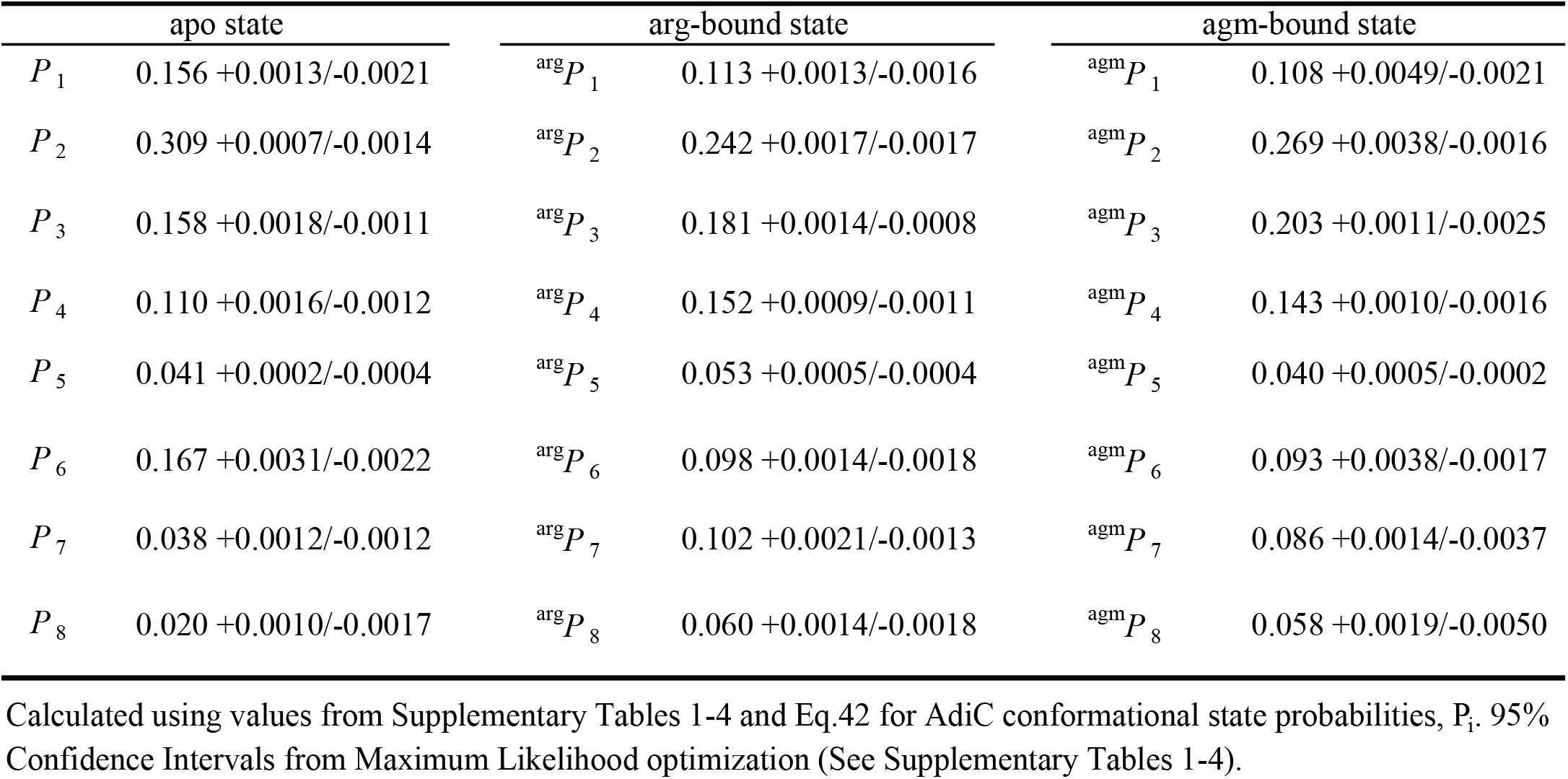
state probabilities in the presence or absence arg or agm

|  | apo state |  | arg-bound state |  | agm-bound state |
| --- | --- | --- | --- | --- | --- |
| $P_1$ | 0.156 +0.0013/-0.0021 | ${}^{\text{arg}}P_1$ | 0.113 +0.0013/-0.0016 | ${}^{\text{agm}}P_1$ | 0.108 +0.0049/-0.0021 |
| $P_2$ | 0.309 +0.0007/-0.0014 | ${}^{\text{arg}}P_2$ | 0.242 +0.0017/-0.0017 | ${}^{\text{agm}}P_2$ | 0.269 +0.0038/-0.0016 |
| $P_3$ | 0.158 +0.0018/-0.0011 | ${}^{\text{arg}}P_3$ | 0.181 +0.0014/-0.0008 | ${}^{\text{agm}}P_3$ | 0.203 +0.0011/-0.0025 |
| $P_4$ | 0.110 +0.0016/-0.0012 | ${}^{\text{arg}}P_4$ | 0.152 +0.0009/-0.0011 | ${}^{\text{agm}}P_4$ | 0.143 +0.0010/-0.0016 |
| $P_5$ | 0.041 +0.0002/-0.0004 | ${}^{\text{arg}}P_5$ | 0.053 +0.0005/-0.0004 | ${}^{\text{agm}}P_5$ | 0.040 +0.0005/-0.0002 |
| $P_6$ | 0.167 +0.0031/-0.0022 | ${}^{\text{arg}}P_6$ | 0.098 +0.0014/-0.0018 | ${}^{\text{agm}}P_6$ | 0.093 +0.0038/-0.0017 |
| $P_7$ | 0.038 +0.0012/-0.0012 | ${}^{\text{arg}}P_7$ | 0.102 +0.0021/-0.0013 | ${}^{\text{agm}}P_7$ | 0.086 +0.0014/-0.0037 |
| $P_8$ | 0.020 +0.0010/-0.0017 | ${}^{\text{arg}}P_8$ | 0.060 +0.0014/-0.0018 | ${}^{\text{agm}}P_8$ | 0.058 +0.0019/-0.0050 |

Calculated using values from Supplementary Tables 1-4 and Eq.42 for AdiC conformational state probabilities, $P_i$ . 95% Confidence Intervals from Maximum Likelihood optimization (See Supplementary Tables 1-4).

**Supplementary Table 6:** Equilibrium dissociation constants for arg or agm bound states

| arg-bound state($\mu\text{M}$) | | agm-bound state($\mu\text{M}$) | |
| --- | --- | --- | --- |
| $^{\text{arg}}K_{\text{D1}}$ | 54 +22/-13 | $^{\text{agm}}K_{\text{D1}}$ | 69 +19/-18 |
| $^{\text{arg}}K_{\text{D2}}$ | 51 +21/-12 | $^{\text{agm}}K_{\text{D2}}$ | 55 +15/-15 |
| $^{\text{arg}}K_{\text{D3}}$ | 34 +15/-7.9 | $^{\text{agm}}K_{\text{D3}}$ | 37 +10/-9.8 |
| $^{\text{arg}}K_{\text{D4}}$ | 29 +12/-6.5 | $^{\text{agm}}K_{\text{D4}}$ | 37 +11/-9.8 |
| $^{\text{arg}}K_{\text{D5}}$ | 31 +13/-7.1 | $^{\text{agm}}K_{\text{D5}}$ | 50 +14/-14 |
| $^{\text{arg}}K_{\text{D6}}$ | 68 +30/-16 | $^{\text{agm}}K_{\text{D6}}$ | 86 +25/-23 |
| $^{\text{arg}}K_{\text{D7}}$ | 15 +6.7/-3.5 | $^{\text{agm}}K_{\text{D7}}$ | 21 +6.4/-5.9 |
| $^{\text{arg}}K_{\text{D8}}$ | 13 +5.5/-3.2 | $^{\text{agm}}K_{\text{D8}}$ | 17 +5.3/-4.6 |

AdiC conformational state probabilities, $P_i$ . 95% Confidence Intervals from Maximum Likelihood optimization (See Supplementary Tables 1-4).

**Supplementary Table 7:** Equilibrium constants in the presence or absence of arg or agm

|  | apo state |  | arg-bound state |  | agm-bound state |
| --- | --- | --- | --- | --- | --- |
| $K_{2,1}$ | 1.98 +0.022/-0.016 | $^{\text{arg}}K_{2,1}$ | 2.13 +0.026/-0.016 | $^{\text{agm}}K_{2,1}$ | 2.48 +0.040/-0.082 |
| $K_{3,1}$ | 1.01 +0.026/-0.016 | $^{\text{arg}}K_{3,1}$ | 1.59 +0.036/-0.024 | $^{\text{agm}}K_{3,1}$ | 1.87 +0.044/-0.101 |
| $K_{4,1}$ | 0.709 +0.020/-0.014 | $^{\text{arg}}K_{4,1}$ | 1.34 +0.025/-0.023 | $^{\text{agm}}K_{4,1}$ | 1.31 +0.033/-0.070 |
| $K_{5,1}$ | 0.265 +0.003/-0.003 | $^{\text{arg}}K_{5,1}$ | 0.463 +0.010/-0.007 | $^{\text{agm}}K_{5,1}$ | 0.367 +0.007/-0.013 |
| $K_{6,1}$ | 1.07 +0.018/-0.007 | $^{\text{arg}}K_{6,1}$ | 0.866 +0.011/-0.012 | $^{\text{agm}}K_{6,1}$ | 0.856 +0.016/-0.015 |
| $K_{7,1}$ | 0.247 +0.013/-0.009 | $^{\text{arg}}K_{7,1}$ | 0.896 +0.031/-0.021 | $^{\text{agm}}K_{7,1}$ | 0.799 +0.029/-0.066 |
| $K_{8,1}$ | 0.131 +0.007/-0.011 | $^{\text{arg}}K_{8,1}$ | 0.530 +0.016/-0.019 | $^{\text{agm}}K_{8,1}$ | 0.535 +0.028/-0.067 |

**Supplementary Table 8:** Observed and calculated k_max_ and K_m_ of Arg and Agm

| **Ligand** | **Arg** |  |  |  | **Agm** |  |  |  |
| --- | --- | --- | --- | --- | --- | --- | --- | --- |
| **Configuration** | Inside-out |  | Outside-out |  | Inside-out |  | Outside-out |  |
| | $k_{\max}$ (s<sup>-1</sup>) | $K_m$ (mM) | $k_{\max}$ (s<sup>-1</sup>) | $K_m$ (mM) | $k_{\max}$ (s<sup>-1</sup>) | $K_m$ (mM) | $k_{\max}$ (s<sup>-1</sup>) | $K_m$ (mM) |
| **Observed<sup>a</sup>** | 0.120 | 0.100 | 0.038 | 0.052 | 0.125 | 0.100 | 0.086 | 0.043 |
| **Calculated<sup>b</sup>** | 0.096 | 0.028 | 0.096 | 0.047 | 0.085 | 0.036 | 0.085 | 0.055 |
|  | +0.013/ | +0.009/ | +0.011/ | +0.019/ | +0.022/ | +0.014/ | +0.019/ | +0.023/ |
|  | -0.014 | -0.008 | -0.015 | -0.015 | -0.027 | -0.010 | -0.037 | -0.019 |

<sup>a</sup> values calculated from studying arginine transport for AdiC in liposomes
<sup>b</sup> values calculated from kinetic model for AdiC observed from single molecule polarization studies

